# Genomes of the entomopathogenic nematode *Steinernema hermaphroditum* and its associated bacteria

**DOI:** 10.1101/2025.01.09.632278

**Authors:** Erich M. Schwarz, Anil Baniya, Jennifer K. Heppert, Hillel T. Schwartz, Chieh-Hsiang Tan, Igor Antoshechkin, Paul W. Sternberg, Heidi Goodrich-Blair, Adler R. Dillman

**Affiliations:** Department of Molecular Biology and Genetics, Cornell University, Biotechnology 407, Ithaca, NY 14853, U.S.A; Department of Nematology, University of California-Riverside, 900 University Ave., Riverside, CA 92521, U.S.A; Department of Microbiology, University of Tennessee-Knoxville, 1311 Cumberland Ave., Knoxville, TN 37996, U.S.A; Division of Biology and Biological Engineering, California Institute of Technology, 1200 E. California Blvd., Pasadena, CA 91125, U.S.A

## Abstract

As an entomopathogenic nematode (EPN), *Steinernema hermaphroditum* parasitizes insect hosts and harbors symbiotic *Xenorhabdus griffinae* bacteria. In contrast to other Steinernematids, *S. hermaphroditum* has hermaphroditic genetics, offering the experimental scope found in *Caenorhabditis elegans*. To enable biological analysis of *S. hermaphroditum*, we have assembled and analyzed its reference genome. This genome assembly has five chromosomal scaffolds and 83 unassigned scaffolds totaling 90.7 Mb, with 19,426 protein-coding genes having a BUSCO completeness of 88.0%. Its autosomes show higher densities of strongly conserved genes in their centers, as in *C. elegans*, but repetitive elements are evenly distributed along all chromosomes, rather than with higher arm densities as in *C. elegans*. Either when comparing protein motif frequencies between nematode species or when analyzing gene family expansions during nematode evolution, we observed two categories of genes preferentially associated with the origin of *Steinernema or S. hermaphroditum:* orthologs of venom genes in *S. carpocapsae* or *S. feltiae*; and some types of chemosensory G protein-coupled receptors, despite the tendency of parasitic nematodes to have reduced numbers of chemosensory genes. Three-quarters of venom orthologs occurred in gene clusters, with the larger clusters comprising functionally diverse pathogenicity islands rather than paralogous repeats of a single venom gene. While assembling the genome of *S. hermaphroditum*, we coassembled bacterial genomes, finding sequence data for not only the known symbiont, *X. griffinae*, but also for eight other bacterial genera. All eight genera have previously been observed to be associated with Steinernema species or the EPN *Heterorhabditis*, and may constitute a “second bacterial circle” of EPNs. The genome assemblies of *S. hermaphroditum* and its associated bacteria will enable use of these organisms as a model system for both entomopathogenicity and symbiosis.

## Introduction

Nematodes are best known for including the model organism *Caenorhabditis elegans* (Corsi et al. 2015), but *C. elegans* is merely one out of roughly 80 million nematode species with greatly diverse morphologies, life cycles, and ecological niches (De Ley 2006; Larsen et al. 2017). Some nematodes have evolved to infect, kill, and feed on insects, with help from symbiotic bacteria (Dillman and Sternberg 2012). Nematodes of this kind are called entomopathogenic, and are valuable for both basic research and applied biology. Scientifically, they provide complex ecological models of both animal parasitism and microbial symbiosis (Goodrich-Blair 2007; Stock 2019; Ogier et al. 2023); commercially, they provide natural non-chemical insecticides (Tarasco et al. 2023).

The two dominant genera of entomopathogenic nematodes (EPNs) are *Heterorhabditis* and *Steinernema*. Genomics of the latter began with the sequencing and analysis of five *Steinernema* species (Dillman et al. 2015; Rougon-Cardoso et al. 2016; Fu et al. 2020), of which *S. carpocapsae* has been the most extensively studied; these studies have included genomics of putative *cis*-regulatory DNA motifs (Dillman et al. 2015), embryonic transcriptomics (Macchietto et al. 2017), neuropeptides and miRNAs potentially regulating behavior (Morris et al. 2017; Warnock et al. 2019; Warnock et al. 2021), transcriptomic responses to symbiotic bacteria (Lefoulon et al. 2022), identification of venom proteins secreted from activated infectious juveniles that are conserved in other *Steinernema* species (Lu et al. 2017; Chang et al. 2019), immunosuppression of insects by venom proteins (Parks et al. 2021; Jones et al. 2022; Parks et al. 2023), and genome reassembly with long-read sequencing data that generated a single X-chromosomal scaffold (Serra et al. 2019).

*Heterorhabditis* and *Steinernema* nematodes symbiotically associate with *Photorhabdus* and *Xenorhabdus* bacteria, respectively; the bacterial symbiotes help kill insect hosts and consume insect tissues, cserve as a food source for the nematode, and generate secondary metabolites that protect the nematodes from other microbes or predators (Goodrich-Blair 2007; Mucci et al. 2022; Puza and Machado 2024). Each *Steinernema* species typically is associated with a specific *Xenorhabdus* species that is carried between insect hosts within an anterior intestinal receptacle (Snyder et al. 2007). Complicating this simple picture is that a more extended set of bacterial species have been found associated with EPNs in nature, intermittently but repeatedly enough that they have been called a “second bacterial circle” or “pathobiome”, and suggested to be commensals or even symbionts of *Steinernema* and other EPNs (Ogier et al. 2023).

Although there are benefits to studying *S. carpocapsae*, it also has limitations as an experimental model for exploring EPN biology. Its most basic limitation is that it is a male-female species, unlike nematodes such as *Caenorhabditis elegans* or *Pristionchus pacificus* that are primarily self-fertilizing hermaphrodites with a small fraction of males. As Brenner recognized five decades ago, self-fertilization facilitates the isolation and characterization of mutations (Brenner 1974); more generally, both classical and reverse genetics can be done more easily in nematode species with the hermaphroditic sexual mode of *C. elegans* (Corsi et al. 2015). Fortunately, the recently reisolated species *Steinernema hermaphroditum* (Griffin et al. 2001; Stock et al. 2004; Bhat et al. 2019) is also a self-fertilizing hermaphrodite with *C. elegans*-like genetics, despite being evolutionarily remote from *C. elegans* (Figure 1). Thus, we and others have begun developing *S. hermaphroditum* as a new model for studying parasitic and mutualistic symbiosis (Cao et al. 2022; Garg et al. 2022; Cao 2023; Huynh et al. 2023; Rodak et al. 2024; Schwartz et al. 2024).

**Figure 1.**
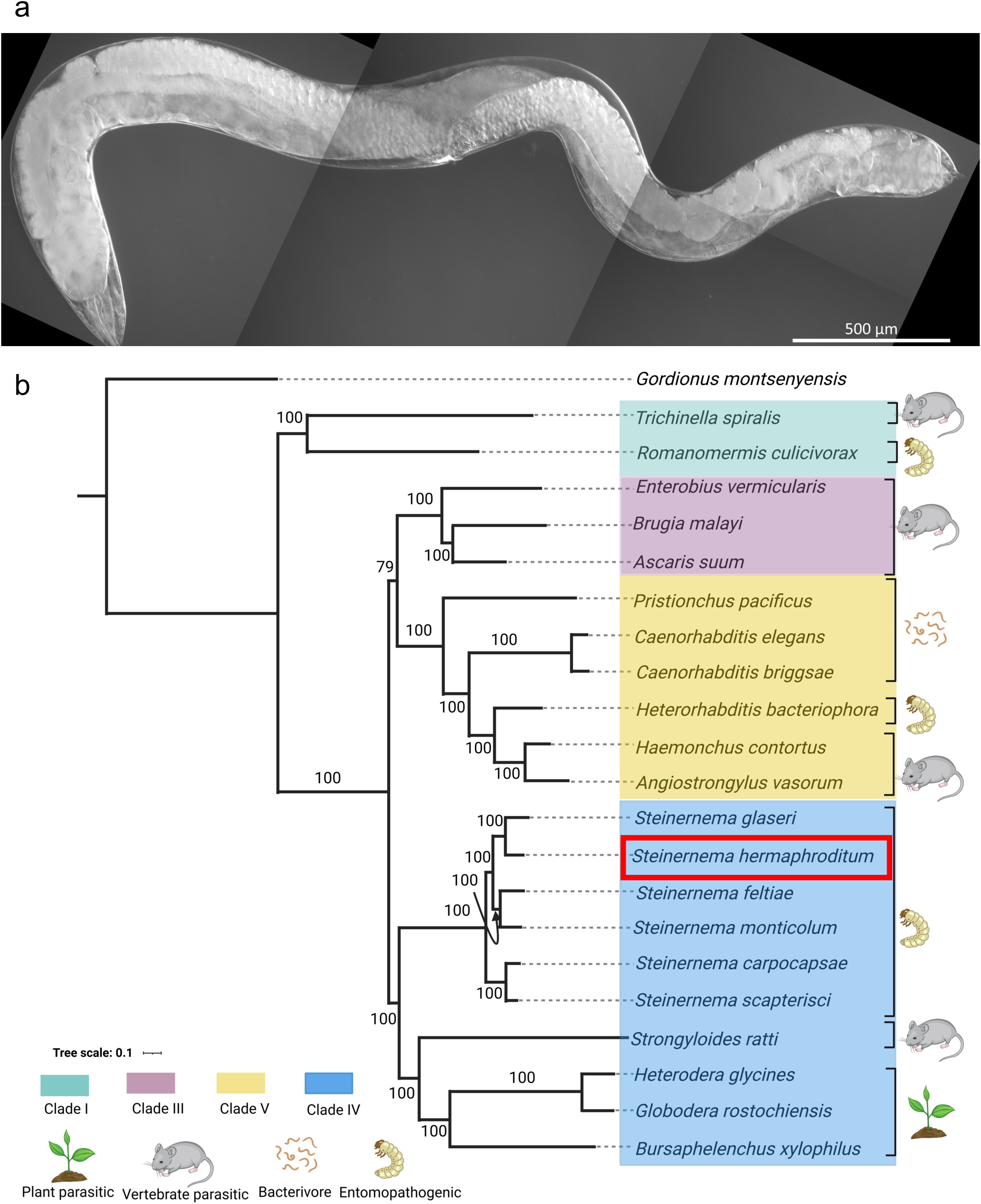
Morphology and phylogeny. (a) An adult *S. hermaphroditum* hermaphrodite, viewed from the side with its anterior pharynx to the left; scale bar, 500 µm. Its body organization resembles that of *C. elegans*, but is roughly four times longer, which fits an observed tendency of parasitic nematodes to be larger than free-living nematodes (Yeates and Boag 2006). The body size of adult *S. hermaphroditum* depends on culture conditions; the adult here was fed on *Comamonas aquatica* bacteria to optimize its growth (Rodak et al. 2024). (b) A phylogeny of *S. hermaphroditum*. This phylogenetic tree relates *S. hermaphroditum* (boxed in red) to a selection of well-studied nematode species spanning clades I, III, IV, and V of the nematode phylum (Kiontke et al. 2021), with the nematomorph *Gordionus montsenyensis* representing an outgroup animal phylum (Rota-Stabelli et al. 2013). Numbers on branching points represent bootstrap values. Ecological niches of the nematodes (plant-parasitic, vertebrate-parasitc, free-living bacteriovore, or entomopathogen) are indicated. Although *S. hermaphroditum* superficially resembles *C. elegans* in its hermaphroditic genetics, the two species are phylogenetically remote (clades IV versus V).

A key resource for any newly developed model organism is an annotated genome assembly of the highest possible quality. We have thus assembled a chromosomal genome sequence for *S. hermaphroditum*. While doing this, from among the coassembled non-nematode contigs we unexpectedly also identified genomes for bacteria resembling the suggested pathobiome of EPNs. Here, we describe and analyze these genomes.

## Results

### *S. hermaphroditum* genome assembly and annotation

To assemble the genome of *S. hermaphroditum* PS9179, we sequenced genomic DNA from adult hermaphrodites (Supplementary Table S1) with long Oxford Nanopore reads (to 137x genome coverage), short Illumina reads (to 106x coverage), and short Omni-C reads (to 392x coverage); we assembled the reads to chromosomal scaffolds with Raven (Vaser and Šikić 2021) and 3D-DNA (Dudchenko et al. 2017). During assembly, we used Racon (Vaser et al. 2017) and POLCA (Zimin and Salzberg 2020) to correct contig sequence errors; we also used sourmash (Pierce et al. 2019), FCS-GX (Astashyn et al. 2024), and OrthoFinder (Emms and Kelly 2019) to detect and remove microbial contigs (analyzed further below). This yielded a final genome assembly with five chromosomal scaffolds totalling 87.9 Mb, along with unscaffolded contigs totalling 2.8 Mb (Table 1). The unscaffolded contigs contained 72% repetitive DNA, which may explain why they could not be linked to chromosomes despite extensive Omni-C data.

**Table 1:** Genome statistics.

|  | *Steinernema hermaphroditum*<br>chromosomes | *Steinernema hermaphroditum*<br>non-chromosomes | *Caenorhabditis elegans* | *Steinernema carpocapsae* | *Strongyloides ratti* |
| --- | --- | --- | --- | --- | --- |
| Total nt | 87,891,628 | 2,836,821 | 100,286,401 | 84,512,609 | 43,166,851 |
| Scaffolds | 5 | 83 | 7 | 16 | 136 |
| Contigs | 62 | 96 | 7 | 32 | 311 |
| % non-N | 99.97 | 99.77 | 100.00 | 99.46 | 99.41 |
| % GC | 47.06 | 45.83 | 35.44 | 45.67 | 21.43 |
| % repeats | 15.94 | 71.54 | 21.95 | 9.78 | 21.86 |
| Scaffold N50 nt | 17,711,000 | 60,131 | 17,493,829 | 7,362,381 | 11,693,564 |
| Contig N50 nt | 5,566,357 | 55,000 | 17,493,829 | 4,209,505 | 1,164,709 |
| Protein-coding genes | 19,275 | 151 | 19,983 | 30,931 | 12,464 |
| Protein isoforms | 35,972 | 293 | 28,591 | 36,703 | 12,484 |
| Genome completeness | 75.4% | 0.5% | 98.7% | 73.8% | 67.5% |
| Proteome completeness | 87.7% | 0.8% | 100.0% | 75.3% | 70.7% |
| Reference | This study | This study | (Consortium 1998) | (Serra et al. 2019) | (Hunt et al. 2016) |

Genome and proteome completeness scores were estimated with BUSCO (Waterhouse et al., 2018). Statistics for published genomes were taken from WormBase ParaSite (Howe et al. 2017).

The genus *Steinernema* is known to have five chromosomes (Curran 1989; Cao et al. 2022), but their syntenic identities were unclear. We used whole-genome alignments and conserved gene sets from vis_ALG to identify Nigon elements (Gonzalez de la Rosa et al. 2021) in the five chromosomal scaffolds of *S. hermaphroditum* (Figure 2a). Three autosomal chromosomes predominantly contained genes from the Nigon elements A, C, and E, corresponding to *C. elegans* chromosomes I, III, and V, and were named accordingly. One autosome matched Nigon element N, with no exact *C. elegans* cognate, and so was named chromosome N. The fifth chromosome was syntenic to the X chromosome of *S. carpocapsae* (Serra et al. 2019), with its central 3 Mb matching Nigon element X, and was thus identified as chromosome X (Figure 2b). Whole-genome alignment of protein-coding sequences between *S. hermaphroditum* and *C. elegans* was consistent with these chromosomal assignments (Figure 2c).

**Figure 2.**
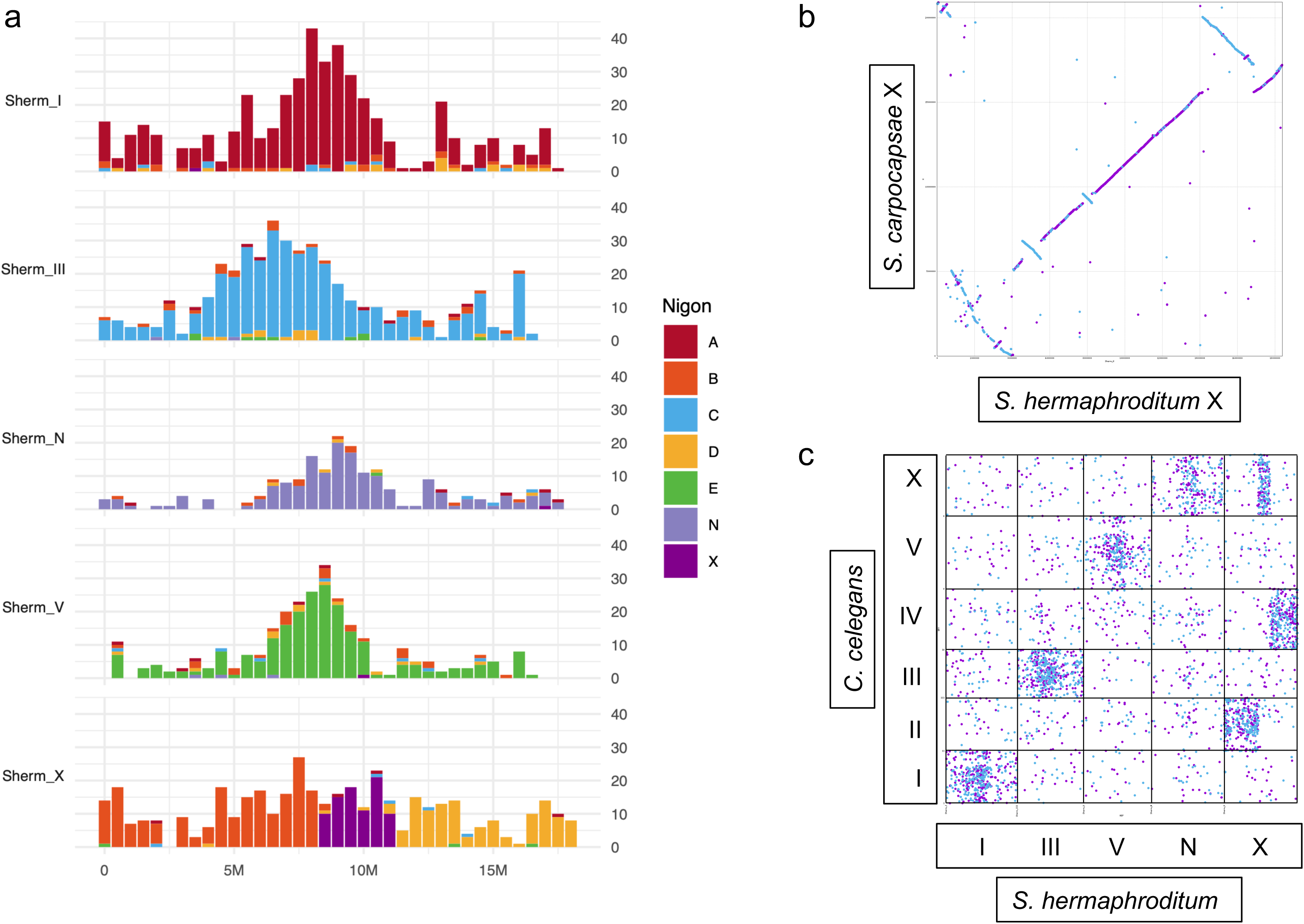
Chromosomal syntenies. (a) Ancestral elements (Nigon elements) of *S. hermaphroditum* chromosomes. These were detected by mapping 2,191 genes previously identified as having strict (1:1) orthologies between most nematode species, and as consistently falling into groups of seven chromosomally colocating loci (Gonzalez de la Rosa et al. 2021). Three Nigon elements (A, C, and E) identify chromosomes I, III, and V as being syntenic with their *C. elegans* counterparts. One chromosome is identified by only element N, with no exact *C. elegans* equivalent. The X chromosome carries three fused elements, B, X, and D; this fits a recurrent pattern of X elements fusing with other elements during nematode X-chromosomal evolution (Gonzalez de la Rosa et al. 2021). (b) Synteny between the X chromosomes of *S. hermaphroditum* and *S. carpocapsae*. The latter’s X-chromosomal status was confirmed by whole-genome sequencing coverage of males versus females, with males having 50% the coverage of females (Serra et al. 2019). Synteny is unbroken from the central region (X element) into the left arm (D element), but is rearranged at the junction of the right arm (B element). (c) Synteny between *S. hermaphroditum* and *C. elegans* chromosomes. Dots represent mutual best matches of protein-coding open reading frames (likely exons). Patterns of synteny are consistent with Nigon element analysis.

To identify protein-coding genes in *S. hermaphroditum*, we generated RNA-seq from mixed-stage hermaphrodites and from infective juveniles, to capture expression from a diverse gene set that would optimally guide gene prediction. We then used both our RNA-seq data and protein sequence data from clade IV nematodes (Supplementary Table S7) with BRAKER2 and TSEBRA (Brůna et al. 2021; Gabriel et al. 2021) to predict 19,426 genes, of which all but 151 could be assigned to chromosomes (Supplementary Table S2; Supplementary Files S3-S5). These genes compared favorably in their BUSCO completeness (88.0%) to two well-characterized gene sets from other species in clade IV, *S. carpocapsae* (75.3%) and *Strongyloides ratti* (70.7%; Table 1) (Hunt et al. 2016; Serra et al. 2019). We identified 1,906 ncRNA-coding genomic loci (Supplementary Files S6-S7) by scanning the *S. hermaphroditum* genome with INFERNAL and Rfam 14.10 (Nawrocki and Eddy 2013; Kalvari et al. 2021). To identify repetitive DNA elements, we scanned the genome with RepeatModeler2 (Flynn et al. 2020), filtered the predictions against known protein-coding or ncRNA nematode genes, and mapped the filtered elements with RepeatMasker; this defined 15.9% of the *S. hermaphroditum* genome as repetitive (Supplementary Files S8-S10).

### Chromosomal distribution of genes and repeats

In nematodes, genes and repetitive elements show varying chromosomal densities, which are thought to arise from varying recombination rates and evolutionary selection (Carlton et al. 2022). In *C. elegans*, chromosomes have roughly equal densities of protein-coding genes throughout their length, but higher densities of strongly conserved genes in their centers and repetitive elements in their arms (Carlton et al. 2022). For protein-coding genes of *S. hermaphroditum*, we observed a chromosomal pattern similar to *C. elegans*, with equal densities of protein-coding genes (Figure 3) but higher densities of strongly conserved genes (defined as marker genes for Nigon elements) in autosomal centers than in arms (Figure 2a). However, our analysis of repetitive DNA elements showed little evidence for higher density in arms (Figure 3) even though we could reproduce that pattern for *C. elegans* itself (Supplementary Figure S1). Instead, we observed a generally even distribution of repetitive elements, punctuated by short blocks of highly repetitive DNA that in some cases divide autosomal centers from arms. Evenly distributed repetitive elements have also been observed in *S. ratti* chromosomes (Hunt et al. 2016), and may be more common in nematodes than previously appreciated.

**Figure 3:**
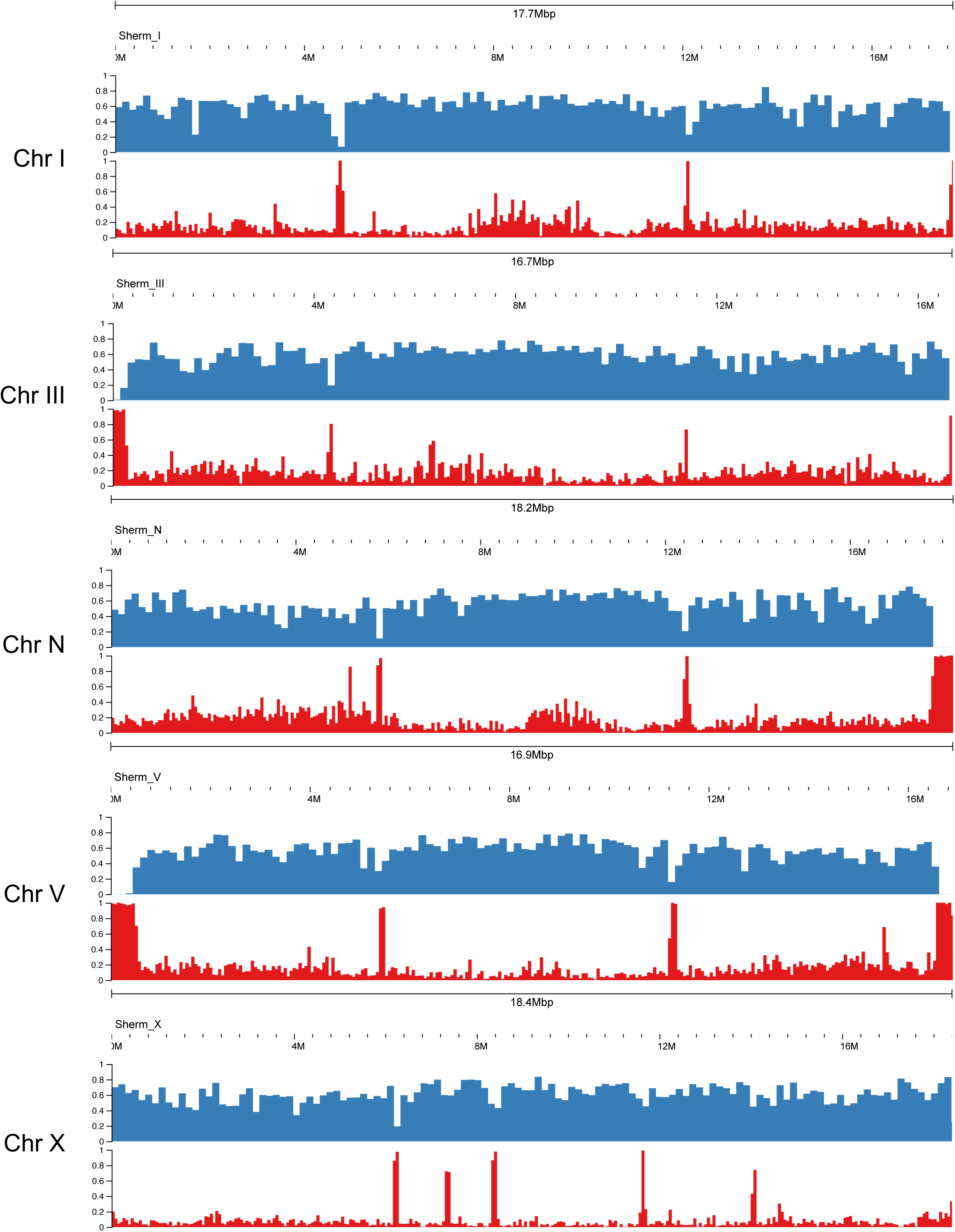
Chromosomal distribution of genes and repeats. Densities of protein-coding genes (blue) and repetitive DNA regions (red) are shown for *S. hermaphroditum* chromosomes. Scale bars on the y-axis indicate the average density per nucleotide, binned for visibility by JBrowse2 (Diesh et al. 2023).

### Protein motifs, functional annotations, and homologies

To identify possible functions of *S. hermaphroditum* protein-coding genes, we computed their protein motifs, Gene Ontology (GO) terms, and orthologies (Supplementary Table S2). We computed 13,170 genes (67.8%) to encode a protein motif from the Pfam or InterPro databases (Blum et al. 2021; Mistry et al. 2021), and 11,952 genes (61.5%) to be associated with a GO term (Carbon et al. 2021). Out of 19,426 *S. hermaphroditum* genes analyzed with OrthoFinder (Emms and Kelly 2019), 5,112 (26.3%) had strict (1:1) orthologies to genes in *C. elegans*; 12,222 (62.9%) had orthology to non-*Steinernema* nematodes generally; 4,970 (25.6%) had orthology only to other *Steinernema* species; and 2,234 (11.5%) were unique to *S. hermaphroditum* (Supplementary Figure S2). These categories of genes may respectively encode proteins that have functional equivalents in *C. elegans*, are generally required for nematode viability, or are adaptations specific to a genus or species.

Differences between nematode species in how many genes encode a type of protein can reflect adaptive differences between the species. For instance, free-living nematode species encode more putative chemoreceptor genes than fully parasitic species, perhaps because free-living nematodes must navigate complex chemical environments while parasites need not (Wheeler et al. 2020; Sural and Hobert 2021). To search for such differences, we compared gene frequencies of Pfam protein motifs between *S. hermaphroditum* and other nematodes (Figure 4; Table 2; Supplementary Figure S2; Supplementary Table S3). Compared to nonparasitic *C. elegans,* the most significantly enriched motif-encoding genes encoded trypsin homologs; other enriched categories included trypsin inhibitor-like cysteine-rich domains (TILs), aspartyl proteases, astacins, and fatty acid/retinoid-binding (FAR) proteins. These are known components of venom proteins in *S. carpocapsae* and *S. feltiae* (Lu et al. 2017; Chang et al. 2019), and their increased abundance in *S. hermaphroditum* likely reflects its parasitic life cycle. Conversely, *C. elegans* had higher frequencies of several types of sensory G-protein-coupled receptors (Str, Srd, Srh, Srj, Sri, Srw, and Srz), and of nuclear hormone receptors with possible chemosensory functions (Sural and Hobert 2021). However, GPCRs of the Srt, Srx, and Srsx types were more common in *S. hermaphroditum*. Moreover, when compared to *S. carpocapsae* the trend reversed, with Srx, Srh, Sri, Srd, Srv, Str, and Srt GPCRs being more abundant in *S. hermaphroditum*. While free-living nematodes do have more complex chemoreceptor gene repertoires than parasitic ones, more subtle trends of parasitic nematodes excelling in some chemoreceptor types can also exist and may be adaptive for their specialized life cycles.

**Figure 4:**
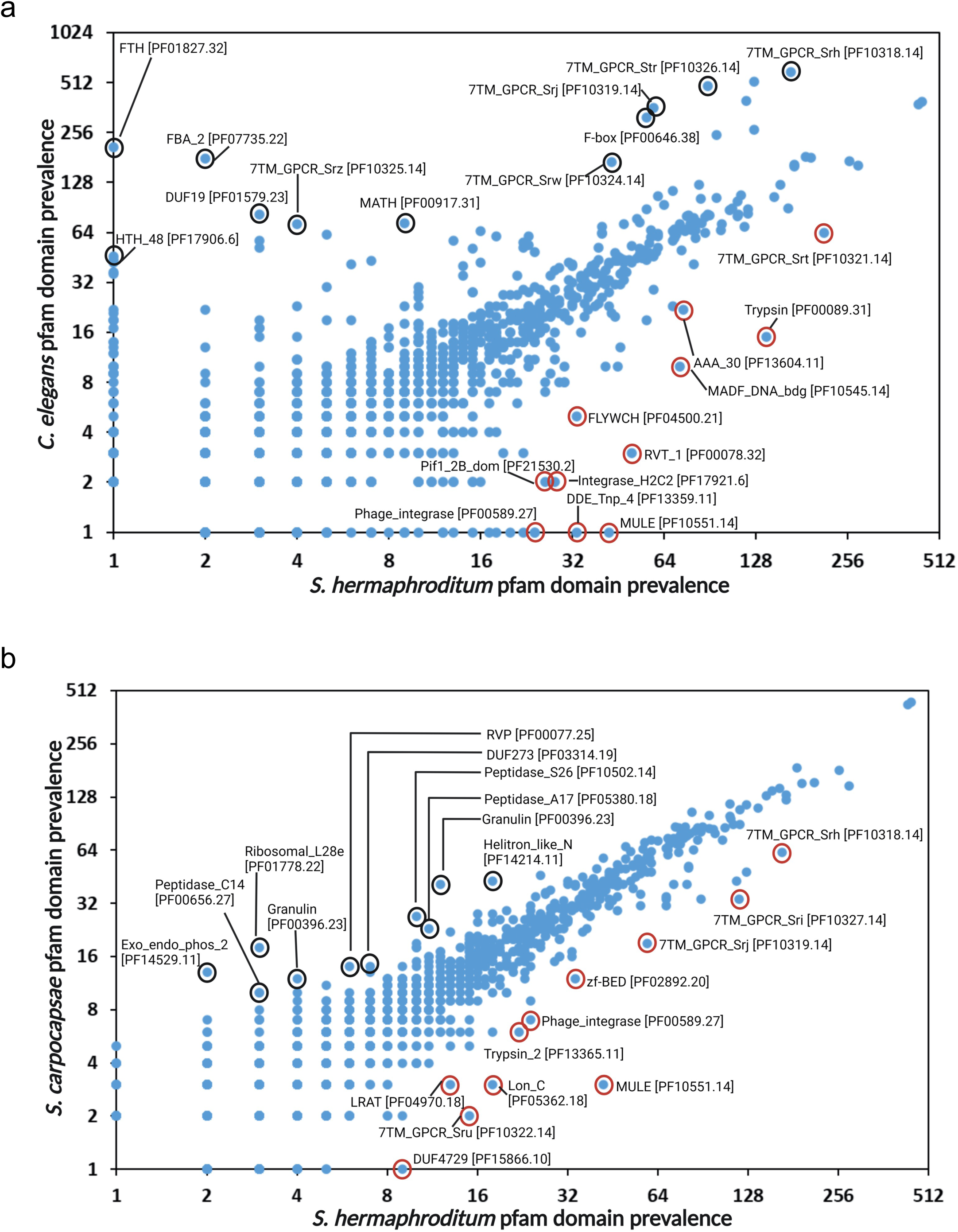
Comparative protein motif frequencies. (a) Comparison of the frequencies of genes encoding Pfam protein motifs between *S. hermaphroditum* and *C. elegans*. (b) Comparison of the frequencies of genes encoding Pfam protein motifs between *S. hermaphroditum* and *S. carpocapsae*. Selected motifs are circled, and are labeled with their Pfam abbreviations and accession numbers.

**Table 2:**
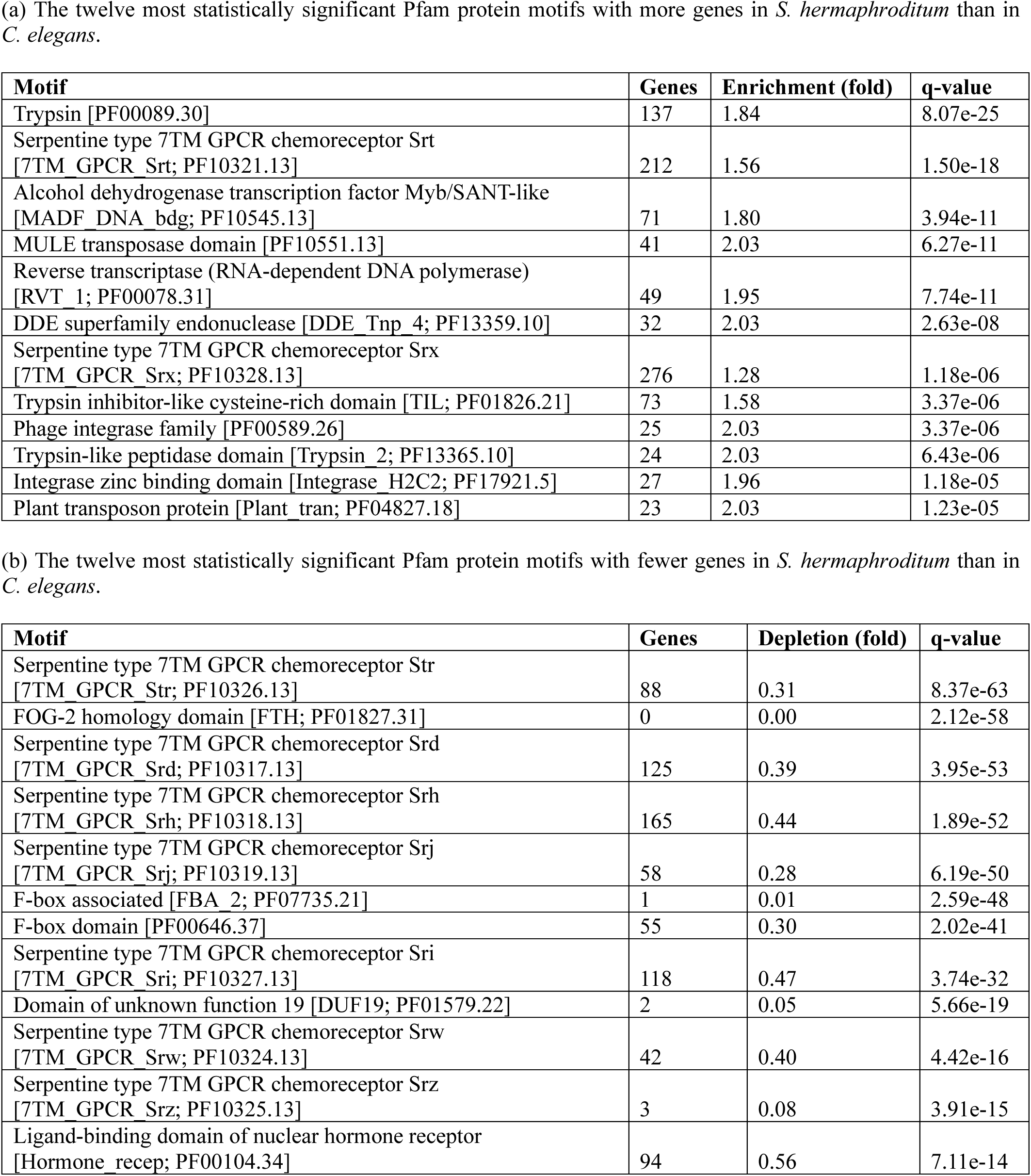
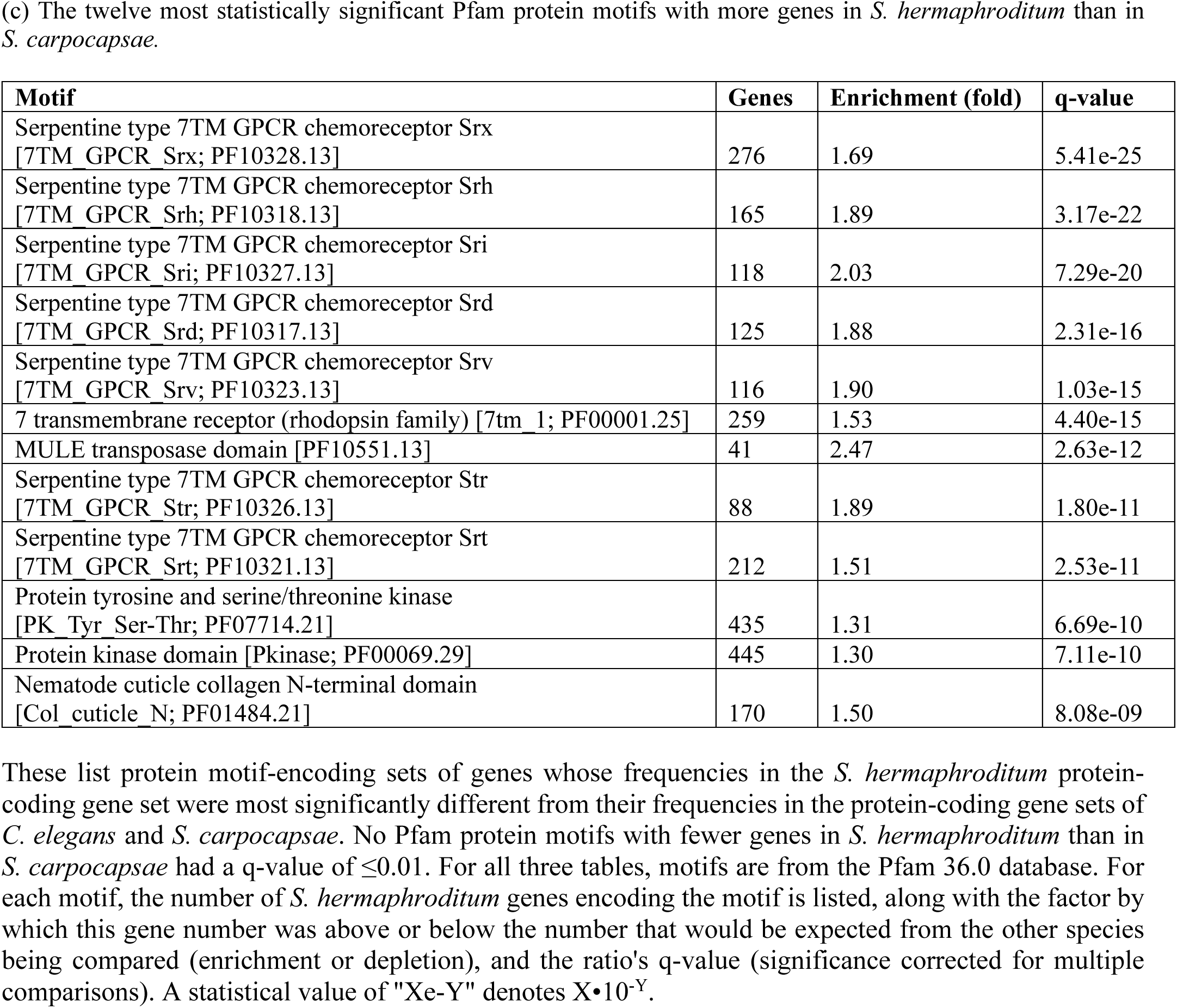
Protein motif-encoding gene comparisons of *S. hermaphroditum*.

### Venom gene orthologs

A key trait of *Steinernema* is that it synthesizes toxic venom proteins and secretes them actively into its hosts (Lu et al. 2017; Chang et al. 2019). Although mass spectrometry has identified 472 venom genes in *S. carpocapsae* and 266 in *S. feltiae*, their chromosomal organization is unknown. To determine this, we first identified 567 genes in *S. hermaphroditum* with orthologies to known venom genes in either *S. carpocapsae* or *S. feltiae*; of these, there were 307 whose orthologs in *S. carpocapsae* or *S. feltiae* were exclusively venom genes (Table 3; Supplementary Table S2). We then used GALEON (Pisarenco et al. 2024) to detect gene clusters of either the more broadly defined 567-gene set or the more narrowly defined or 307-gene set. For the 567 venom orthologs, we found that 432 (76.2%) fell into 82 gene clusters in the genome (Figure 5); for the 307 venom orthologs, we found that 173 (56.4%) fell into 45 gene clusters (Supplementary Figure S3; Supplementary Table S2). Most of the 82 venom ortholog clusters were small (median of three genes, minimum of two genes), but the ten largest clusters comprised nine to 47 genes.

**Figure 5:**
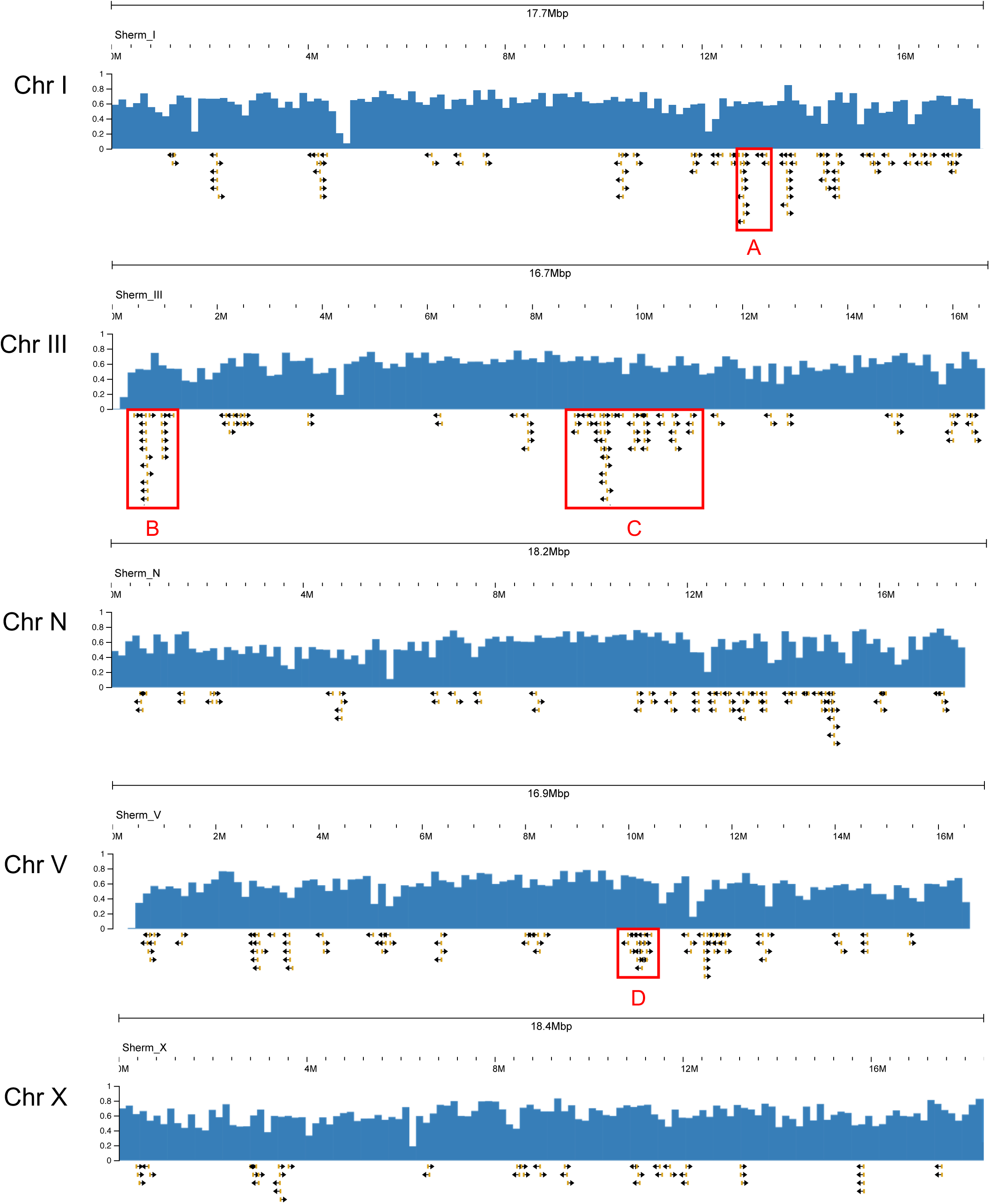
Venom ortholog gene clusters. For each chromosome, the following are shown: mean gene density, and venom ortholog gene clusters identified by GALEON (Pisarenco et al. 2024) from a broadly defined set of 565 venom ortholog genes. Note that these clusters are not due to local increases in gene density, which is essentially uniform. The four largest clusters (Table 4) are marked in red: (A) chromosome I, 11th cluster, with 14 genes; (B) chromosome III, 1st cluster, with 24 genes; (C) chromosome III, 6th cluster, with 47 genes; and (D) chromosome V, 10th cluster with 18 genes.

**Table 3:** Venom gene orthologs in *S. hermaphroditum*.

| **Orthology** | **All orthologs** | **Unique orthologs** |
| --- | --- | --- |
| Either *S. carpocapsae* or *S. feltiae* venom | 567 | 202 |
| *S. carpocapsae* venom | 425 | 160 |
| *S. feltiae* venom | 267 | 71 |
| *S. carpocapsae* and *S. feltiae* venom | 125 | 29 |
| In either *S. carpocapsae* or *S. feltiae*, only venom | 307 | 172 |
| In *S. carpocapsae*, only venom | 244 | 139 |
| In *S. feltiae*, only venom | 93 | 47 |
| In both *S. carpocapsae* and *S. feltiae*, only venom | 30 | 14 |

**Table 4:** Instances of functionally diverse protein motifs overrepresented in venom ortholog gene clusters.

Chromosome III, 6th cluster, with 47 genes:
| Motif | Motif genes | Motif/cluster genes | Enrichment (fold) | q-value |
| --- | --- | --- | --- | --- |
| Trypsin [PF00089.30] | 137 | 14 | 42.2 | 5.07e-16 |
| Nematode fatty acid retinoid binding protein [Gp-FAR-1; PF05823.16] | 40 | 7 | 72.3 | 1.27e-08 |
| Potassium channel inhibitor ShK domain-like [PF01549.28] | 113 | 6 | 21.9 | 0.000474 |
| Astacin [PF01400.28] | 81 | 5 | 25.5 | 0.00176 |
| Peptidase C1-like [PF03051.19] | 13 | 3 | 95.4 | 0.00354 |

Chromosome III, 1st cluster, with 24 genes:
| Motif | Motif genes | Motif/cluster genes | Enrichment (fold) | q-value |
| --- | --- | --- | --- | --- |
| Trypsin [PF00089.30] | 137 | 17 | 100.4 | 1.49e-28 |
| Eukaryotic aspartyl protease [Asp; PF00026.27] | 55 | 4 | 58.9 | 0.000927 |

Chromosome V, 10th cluster, with 18 genes:
| Motif | Motif genes | Motif/cluster genes | Enrichment (fold) | q-value |
| --- | --- | --- | --- | --- |
| TATA box binding protein associated factor [TAF; PF02969.21] | 9 | 4 | 479.7 | 3.08e-07 |
| Lipase (class 3) [PF01764.29] | 30 | 4 | 143.9 | 2.20e-05 |
| Glycosyl hydrolases family 35 [Glyco hydro 35; PF01301.23] | 3 | 2 | 719.6 | 0.00165 |
| Alpha/beta hydrolase fold [Abhydrolase 1; PF00561.24] | 48 | 3 | 67.5 | 0.00668 |

Chromosome I, 11th cluster, with 14 genes:
| Motif | Motif genes | Motif/cluster genes | Enrichment (fold) | q-value |
| --- | --- | --- | --- | --- |
| Potassium channel inhibitor ShK domain-like [PF01549.28] | 113 | 9 | 110.5 | 5.11e-14 |
| Peptidase family M13 [PF01431.25] | 41 | 3 | 101.5 | 0.00494 |

To see whether venom ortholog clusters encoded functionally or evolutionarily related genes, we compared the sets of all protein-coding genes encoding Pfam protein motifs to genes in venom clusters, while also comparing the sets of all genes belonging to orthology groups to genes in venom clusters. In both comparisons, we found overlaps significantly enriched beyond the expected random genomic background. Of the 81 clusters, 30 included genes enriched for both particular protein motifs and particular orthology groups; three clusters were enriched for protein motifs only, and 11 clusters were enriched for orthology groups only (Supplementary Table S4). Collectively, the 173 venom ortholog cluster genes were overrepresented for 55 protein motifs and 47 orthology groups. Each of the four largest venom gene clusters were enriched for two or more protein motifs encoding different functions (Table 4). Thus, venom clusters do not arise from local variations of gene density, and are only partially explained by local duplications of a single venom gene; larger clusters harbor different types of venom genes, and can be considered pathogenicity islands in the *S. hermaphroditum* genome.

### Chronology and gene family changes of the Steinernema clade

We used phylogenomics to estimate the timing and gene family changes associated with the emergence of *Steinernema* and the divergence of *S. hermaphroditum*. To estimate timing, we generated a phylogeny for *Steinernema* species and representative nematodes of other clades with IQ-TREE (Minh et al. 2020), and estimated divergence times for the tree’s nodes with PAML (Yang and Rannala 2006; Rannala and Yang 2007) (Figure 6). The split of *Steinernema* from all other clade IV species (e.g., *S. ratti* or *P. redivivus*) was the deepest within clade IV, estimated at ∼200 Mya. In contrast, splitting within the *Steinernema* genus was estimated at ∼60 Mya, and *S. hermaphroditum*’s split from other species at ∼30 Mya. By comparison, we estimated divergence times of only ∼40 Mya between different genera of clade IV (*Heterodera* and *Globodera*) and ∼55 Mya between different genera of clade V (*Angiostrongylus* and *Haemonchus*). Over 80 species have been identified as *Steinernema* (Spiridonov and Subbotin 2016); this, coupled with the older origin of the *Steinernema* genus relative to multigenus groups in clades IV and V, suggests it may encompass more biological diversity than other nematode genera. The divergence time of *S. hermaphroditum* from other *Steinernema* is comparable to that estimated for *C. elegans* and *C. briggsae* (∼27 Mya); however, including a closer relative of *S. hermaphroditum* than *S. feltiae* (such as *S. adamsi*) would certainly give a much more recent time (Baniya et al. 2024).

**Figure 6:**
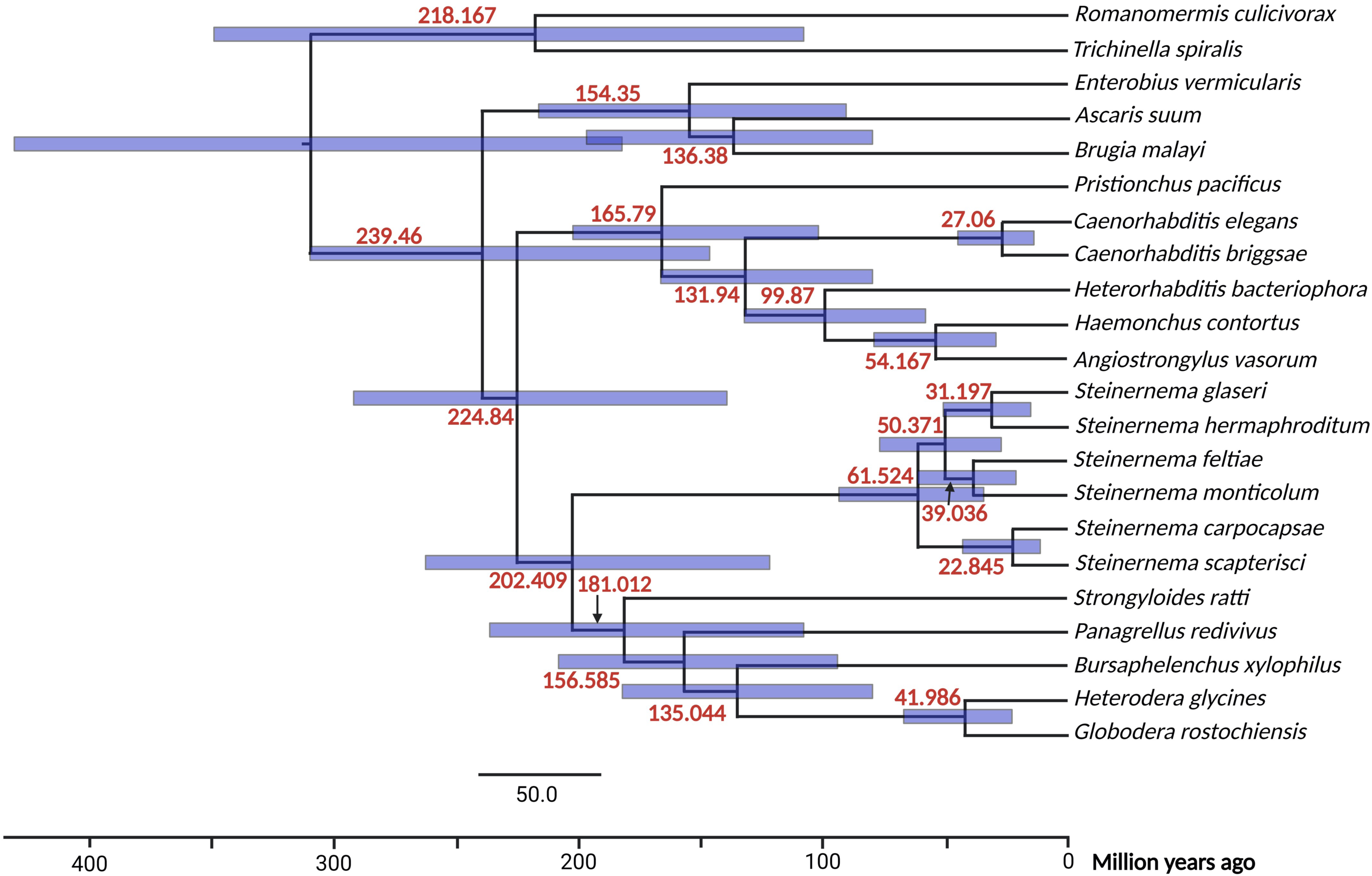
Phylogenetic divergence times. A phylogeny of *S. hermaphroditum* and representative nematode species with estimated divergence times at branch points. Blue error bars indicate 95% confidence intervals. Divergence times and intervals were estimated with PAML (Yang and Rannala 2006; Rannala and Yang 2007).

Changes of gene family size (either through gain or loss of genes) can indicate genetic innovations important for the evolution of a genus or species (Domazet-Loso et al. 2024). To search for such innovations, we estimated which gene families expanded or contracted at the origins of *Steinernema* or *S. hermaphroditum*. We identified gene families as orthology groups of 16 nematode protein-coding gene sets, computed a phylogenetic tree for these 16 taxa, and used CAFE to identify which orthology groups gained or lost gene members at nodes of the tree with a significance of p ≤ 0.05 (Figure 7). At the origin of *Steinernema*, 231 orthology groups expanded significantly; of these, 31 also encoded venom orthologs, and 16 were overrepresented in venom gene clusters. At the splitting off of *S. hermaphroditum*, 159 orthology groups expanded significantly, of which ten also encoded venom orthologs and six were overrepresented in venom gene clusters. No venom gene families significantly contracted at the origin of *Steinernema*; though six contracted significantly at the divergence of *S. hermaphroditum*, none belonged to gene clusters.

**Figure 7:**
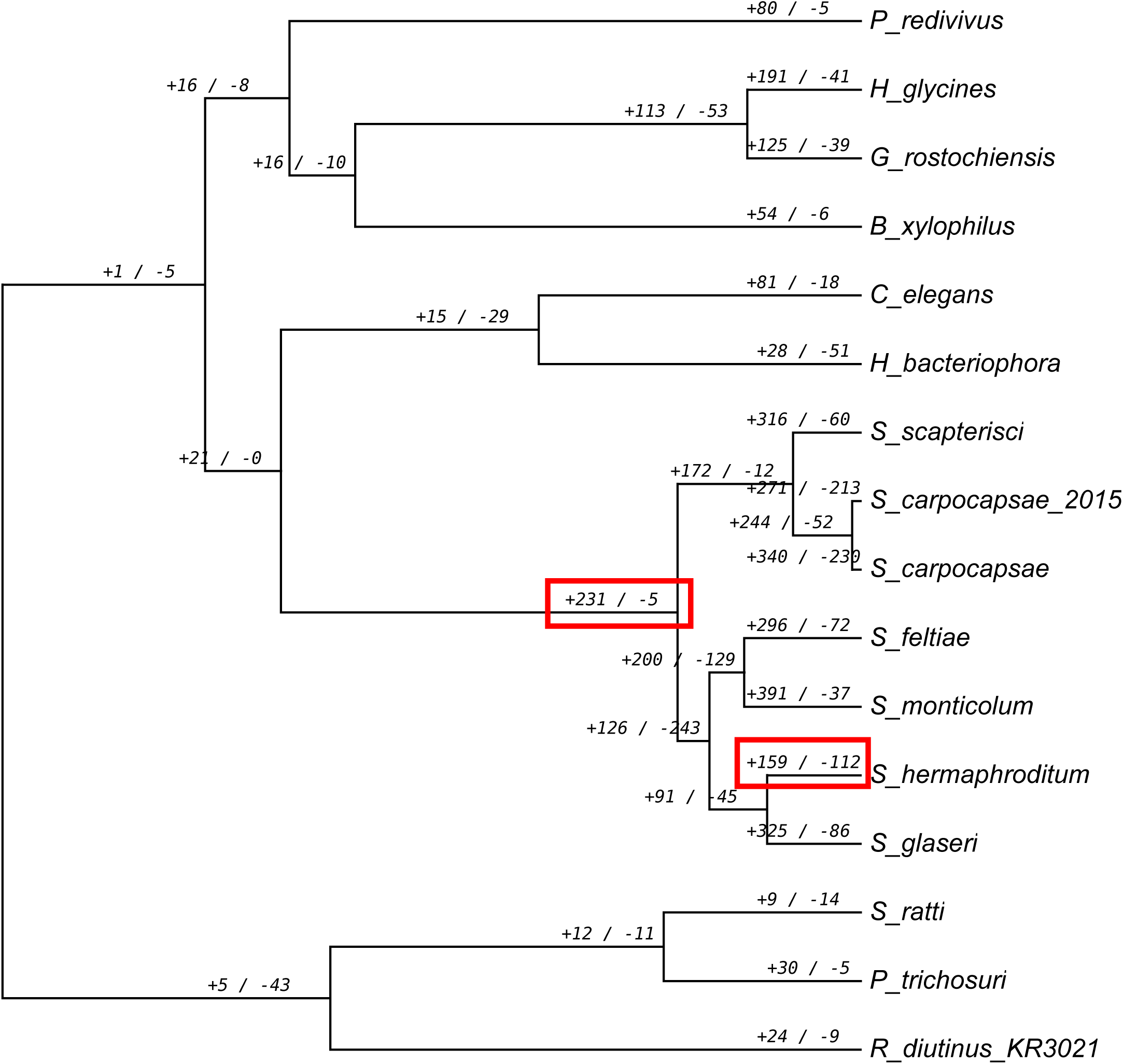
Gene families expanding or contracting in *Steinernema* or *S. hermaphroditum*. Along a phylogenetic tree of 16 nematode protein-coding gene sets, this shows gene families that gained or lost genes at nodes of the tree with a significance of p ≤ 0.05. Gene families were orthology groups computed with OrthoFinder (Emms and Kelly 2019); gains and losses of genes were identified and statistically analyzed with CAFE (Mendes et al. 2021). Key nodes of interest (boxed in red) are the origin of the *Steinernema* genus and the splitting off of *S. hermaphroditum* from other *Steinernema* species. To include known venom genes from *S. carpocapsae* in orthology groups, we analyzed a *S. carpocapsae* gene set from 2015 in addition to the current 2019 version. Note that many other gene families expanded or contracted at nodes of the tree without being scored as statistically significant; for the full set, see Supplementary Figure S4.

Analyzing expanding or contracting gene families more generally for overrepresented protein motifs, we focused on the statistically strongest ones with p ≤ 0.001 (Table 5). At the origin of *Steinernema*, expanding gene families were most enriched for chemosensory receptors (of the Srsx, Srx, Sre, and Srt families), venom-associated protein types (trypsins, trypsin inhibitor-like cysteine-rich domains, and tissue inhibitors of metalloproteinase), and ecdysteroid kinase-like enzymes associated with detoxification (Scanlan et al. 2022). Since very few gene families shrank at the origin of *Steinernema*, no motifs achieved statistical significance. At the splitting off of *S. hermaphroditum*, expanding gene families were, again, most enriched for chemosensory receptors and trypsin, but also for protein motifs associated with mobile DNA elements; contracting gene families of *S. hermaphroditum* were most enriched for a variety of functions, including p450 detoxification enzymes. Results for gene families with p ≤ 0.05 or p ≤ 0.01 were similar to those with p ≤ 0.001 (Supplementary Table S5).

**Table 5:**
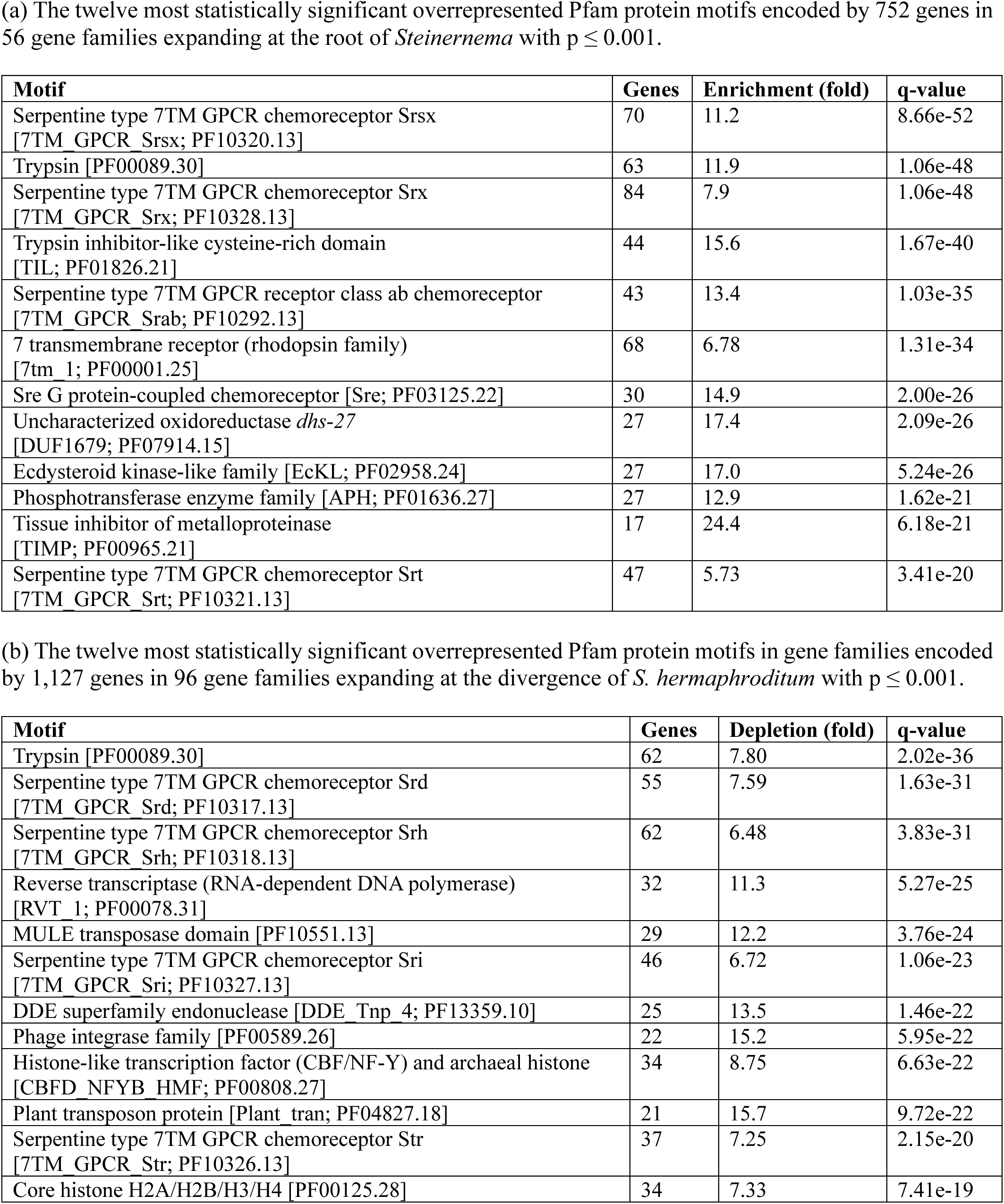
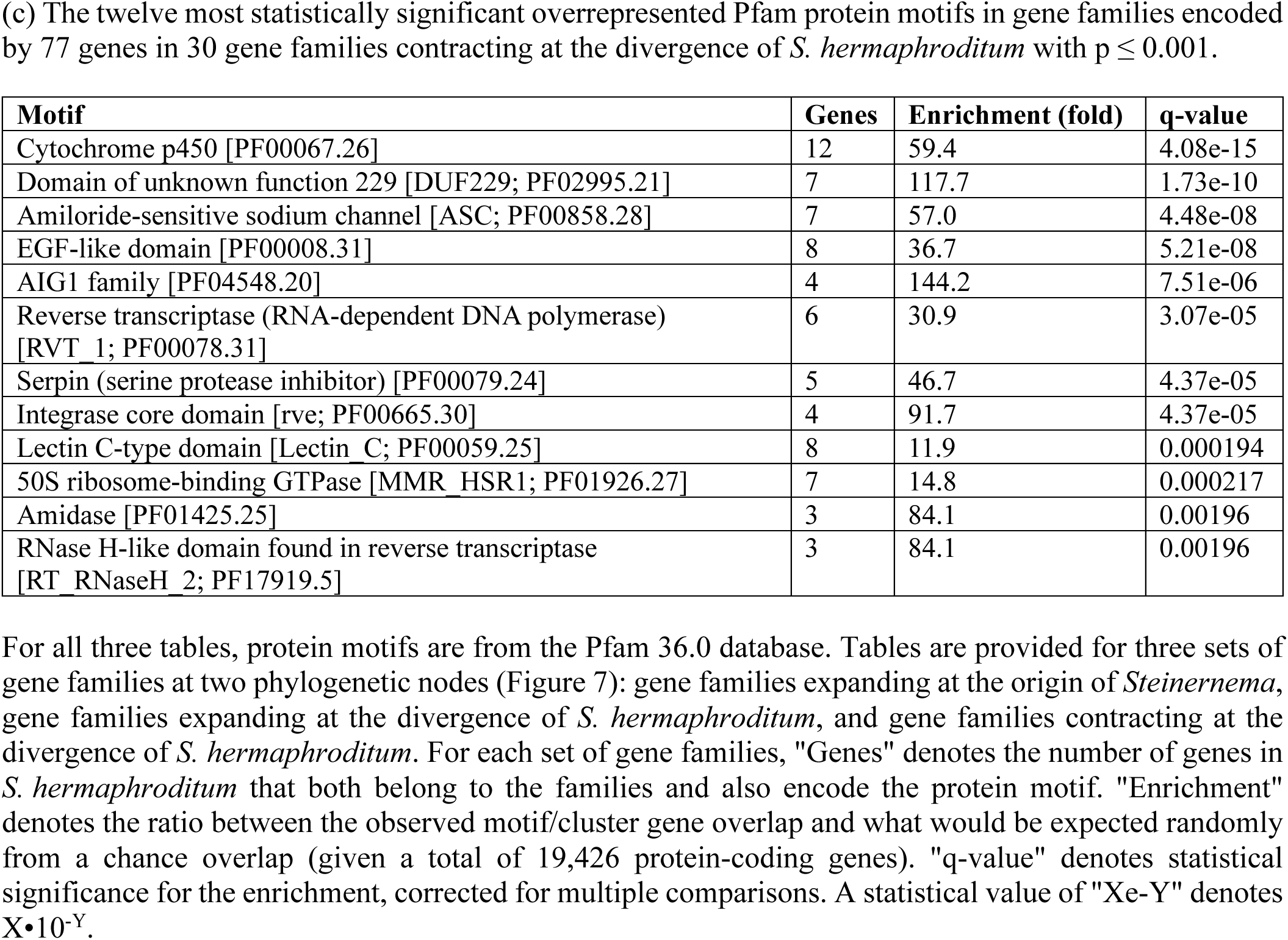
Protein motifs overrepresented in gene families expanding or contracting during *Steinernema* evolution.

| Motif | Genes | Enrichment (fold) | q-value |
| --- | --- | --- | --- |
| Serpentine type 7TM GPCR chemoreceptor Srsx<br>[7TM_GPCR_Srsx; PF10320.13] | 70 | 11.2 | 8.66e-52 |
| Trypsin [PF00089.30] | 63 | 11.9 | 1.06e-48 |
| Serpentine type 7TM GPCR chemoreceptor Srx<br>[7TM_GPCR_Srx; PF10328.13] | 84 | 7.9 | 1.06e-48 |
| Trypsin inhibitor-like cysteine-rich domain<br>[TIL; PF01826.21] | 44 | 15.6 | 1.67e-40 |
| Serpentine type 7TM GPCR receptor class ab chemoreceptor<br>[7TM_GPCR_Srab; PF10292.13] | 43 | 13.4 | 1.03e-35 |
| 7 transmembrane receptor (rhodopsin family)<br>[7tm_1; PF00001.25] | 68 | 6.78 | 1.31e-34 |
| Sre G protein-coupled chemoreceptor [Sre; PF03125.22] | 30 | 14.9 | 2.00e-26 |
| Uncharacterized oxidoreductase *dhs-27*<br>[DUF1679; PF07914.15] | 27 | 17.4 | 2.09e-26 |
| Ecdysteroid kinase-like family [EcKL; PF02958.24] | 27 | 17.0 | 5.24e-26 |
| Phosphotransferase enzyme family [APH; PF01636.27] | 27 | 12.9 | 1.62e-21 |
| Tissue inhibitor of metalloproteinase<br>[TIMP; PF00965.21] | 17 | 24.4 | 6.18e-21 |
| Serpentine type 7TM GPCR chemoreceptor Srt<br>[7TM_GPCR_Srt; PF10321.13] | 47 | 5.73 | 3.41e-20 |

| Motif | Genes | Depletion (fold) | q-value |
| --- | --- | --- | --- |
| Trypsin [PF00089.30] | 62 | 7.80 | 2.02e-36 |
| Serpentine type 7TM GPCR chemoreceptor Srd<br>[7TM_GPCR_Srd; PF10317.13] | 55 | 7.59 | 1.63e-31 |
| Serpentine type 7TM GPCR chemoreceptor Srh<br>[7TM_GPCR_Srh; PF10318.13] | 62 | 6.48 | 3.83e-31 |
| Reverse transcriptase (RNA-dependent DNA polymerase)<br>[RVT_1; PF00078.31] | 32 | 11.3 | 5.27e-25 |
| MULE transposase domain [PF10551.13] | 29 | 12.2 | 3.76e-24 |
| Serpentine type 7TM GPCR chemoreceptor Sri<br>[7TM_GPCR_Sri; PF10327.13] | 46 | 6.72 | 1.06e-23 |
| DDE superfamily endonuclease [DDE_Tnp_4; PF13359.10] | 25 | 13.5 | 1.46e-22 |
| Phage integrase family [PF00589.26] | 22 | 15.2 | 5.95e-22 |
| Histone-like transcription factor (CBF/NF-Y) and archaeal histone<br>[CBFD_NFYB_HMF; PF00808.27] | 34 | 8.75 | 6.63e-22 |
| Plant transposon protein [Plant_tran; PF04827.18] | 21 | 15.7 | 9.72e-22 |
| Serpentine type 7TM GPCR chemoreceptor Str<br>[7TM_GPCR_Str; PF10326.13] | 37 | 7.25 | 2.15e-20 |
| Core histone H2A/H2B/H3/H4 [PF00125.28] | 34 | 7.33 | 7.41e-19 |
| Cytochrome p450 [PF00067.26] | 12 | 59.4 | 4.08e-15 |
| Domain of unknown function 229 [DUF229; PF02995.21] | 7 | 117.7 | 1.73e-10 |
| Amiloride-sensitive sodium channel [ASC; PF00858.28] | 7 | 57.0 | 4.48e-08 |
| EGF-like domain [PF00008.31] | 8 | 36.7 | 5.21e-08 |
| AIG1 family [PF04548.20] | 4 | 144.2 | 7.51e-06 |
| Reverse transcriptase (RNA-dependent DNA polymerase) [RVT 1; PF00078.31] | 6 | 30.9 | 3.07e-05 |
| Serpin (serine protease inhibitor) [PF00079.24] | 5 | 46.7 | 4.37e-05 |
| Integrase core domain [rve; PF00665.30] | 4 | 91.7 | 4.37e-05 |
| Lectin C-type domain [Lectin C; PF00059.25] | 8 | 11.9 | 0.000194 |
| 50S ribosome-binding GTPase [MMR HSR1; PF01926.27] | 7 | 14.8 | 0.000217 |
| Amidase [PF01425.25] | 3 | 84.1 | 0.00196 |
| RNase H-like domain found in reverse transcriptase [RT RNaseH 2; PF17919.5] | 3 | 84.1 | 0.00196 |

### Associated bacterial genomes

Because genomic DNA isolated from nematodes often contains both nematode and microbial contaminant sequences (Kumar et al. 2013; Percudani 2013; Fierst et al. 2017), we used sourmash (Pierce et al. 2019) to exclude non-nematode contigs from the *S. hermaphroditum* genome assembly. This excluded 38 contigs totaling 48,194,651 nt, the largest contigs of which ranged from 1-7 Mb. Usually, microbial contigs found during nematode genome assemblies are discarded and not analyzed further. However, the biology of *S. hermaphroditum* made those contigs potentially interesting, not only because they were likely to include a native symbiont genome, but because *Steinernema* is also known to associate with other bacteria that have not been well characterized, and that may contribute to or detract from its effectiveness against its insect prey (Ogier et al. 2023). We therefore taxonomically classified the 15 largest microbial contigs: 14 of the 15 contigs were bacterial, with the 15th contig not mapping to any known sequence within the Genome Taxonomy Database (Parks et al. 2022) (Figure 8).

**Figure 8.**
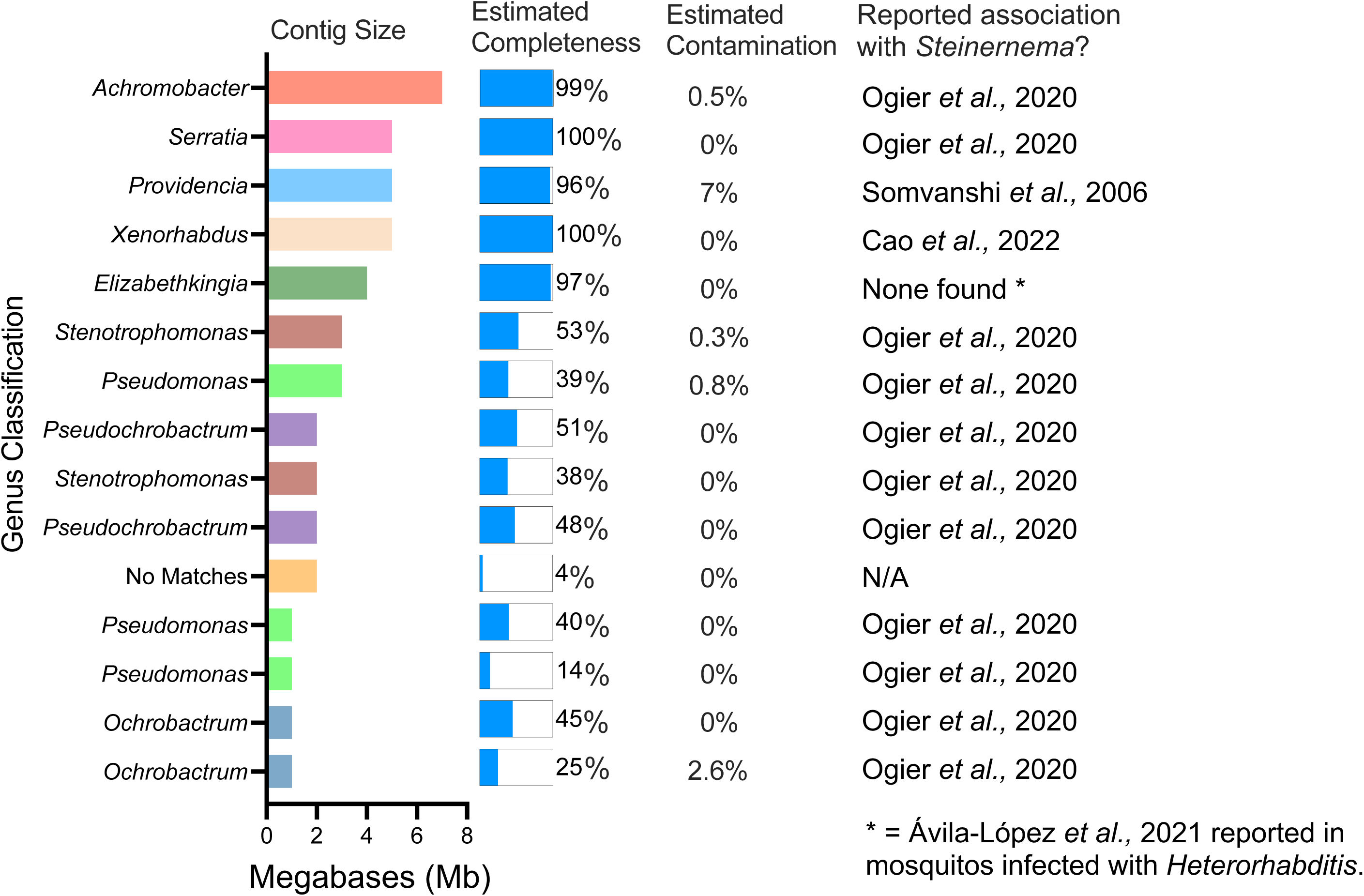
Microbial genomes associated with *S. hermaphroditum*. Taxonomic classification and genetic completeness for the 15 largest microbial contigs coassembled with *S. hermaphroditum*. Estimated contamination levels and previously observed associations with *Steinernema* or other EPNs are noted. Taxonomy and completeness were determined with GTDB-Tk and CheckM (Parks et al. 2015; Chaumeil et al. 2019).

Of the 15 largest contigs, 12 were from microbial genera previously identified as being associated with *Steinernema* (Somvanshi et al. 2006; Ogier et al. 2020; Ávila-López et al. 2021; Ogier et al. 2023) (Figure 8). The five largest contigs represented nearly complete bacterial genomes, according to CheckM (Parks et al. 2015). One of these was from the primary symbiont, *Xenorhabdus griffiniae* HGB2511, which had been used to propagate the nematodes prior to sequencing (Cao et al. 2022). Of the other four largest contigs, three represented bacterial genera previously observed with *Steinernema*: *Achromobacter*, *Providencia*, and *Serratia* (Somvanshi et al. 2006; Ogier et al. 2020). A fourth contig represented *Elizabethkingia*; although not observed with *Steinernema*, this genus was observed with the EPN *Heterorhabditis indica* (Ávila-López et al. 2021). Nine other, smaller contigs were more fragmentary; but when concatenated and tested by CheckM, they also represented largely complete genomes of other bacteria seen with *Steinernema* (Ogier et al. 2020): 99% of a single *Pseudomonas* genome, 97% of a single *Pseudochrobactrum* genome, 97% of a single *Stenotrophomonas* genome, and 66% of a single *Ochrobactrum* genome. These microbial genome assemblies not only included the known symbiont of *S. hermaphroditum*, but also unexpectedly captured eight bacterial species that may represent natural, recurring commensals of *Steinernema*.

## Discussion

To enable its use as a new model organism for both insect parasitism and microbial symbiosis, we have generated and characterized a reference genome for *S. hermaphroditum*. We statistically analyzed differentially represented protein motifs and phylogenetically expanding orthology groups to search for functions particularly associated with the evolution of *Steinernema* as an EPN. We also used the contiguity of our genome to search for chromosomal regions associated with venom gene orthologs. We found that venom genes of *S. hermaphroditum* have four strong (though not universal) tendencies: they encode protein motifs disproportionately represented in *S. hermaphroditum* versus the free-living model *C. elegans*; they belong to protein-coding gene families that expanded at the origin of *Steinernema*, and in some instances kept expanding with the divergence of *S. hermaphroditum*; they occupy gene clusters rather than being evenly distributed along chromosomes; and the larger venom gene clusters are multifunctional, encoding two or more sets of genes encoding nonhomologous proteins related only by their biological role in killing insect hosts.

Having a chromosomal assembly with a well-annotated protein-coding gene set let us test whether venom ortholog genes likely to be associated with virulence fell into discrete genomic regions or not. Genes associated with virulence are not peculiar to *Steinernema*; many nematodes infecting either animals or plants secrete proteins into their hosts that enable parasitism (Abuzeid et al. 2020; Molloy et al. 2024). However, until third-generation genome assembly became routine, most parasitic nematode genome assemblies were fragmented drafts based on short-read sequencing (Doyle 2022); therefore, the batteries of parasitic nematode genes associated with virulence could not be checked for chromosomal organization. We predict that genomic pathogenicity islands of the kind we describe here for *S. hermaphroditum* may be commonplace among parasitic nematodes. One hint that this may be true is that chromosomal-quality genome assemblies for *Heligmosomoides bakeri*, *H. polygyrus*, and *Strongyloides ratti* (intestinal parasites of rodents) have regions of elevated genomic diversity disproportionately populated by genes encoding possible effectors of parasitism (Cole et al. 2023; Stevens et al. 2023). Virulence gene clusters in parasitic nematodes are more likely to encode paralogs sharing a single biochemical function and to be regulated by common signals during the parasitic life cycle; understanding them should thus enable better control of parasitic effectors, whether the goal is to engineer stronger virulence (as is biotechnologically desirable for EPNs) or to block virulence (as is biomedically desirable for animal- or plant-parasitic nematodes).

In addition to venom genes, another category of genes associated with *Steinernema* evolution were classes of chemosensory GPCRs that seem to have become more prevalent as *S. hermaphroditum* evolved. This was unexpected, because recent analyses indicate that known or suspected chemosensory genes encoding either GPCRs or nuclear hormone receptor (NHR) homologs are less abundant in parasitic nematodes than in free-living ones (Wheeler et al. 2020; Sural and Hobert 2021). This is an evolutionarily plausible trend; as nematodes became increasingly parasitic, they might well need less chemosensory navigation, and thus might tend to lose newly superfluous chemoreceptor genes. It is thus unsurprising that several types of GPCRs, along with NHRs, proved more abundant in *C. elegans* than *S. hermaphroditum*; however, it was surprising to observe some types of GPCRs not only were more abundant in *S. hermaphroditum* but also belonged to gene families likely to have expanded at the origin of *Steinernema*. Here, too, our analysis could easily be repeated with many other parasitic nematodes, and might also show that their chemosensory genes have evolved in a more complicated way than previously thought: although parasitic nematodes have fewer chemosensory genes overall, their lifestyle may require increased numbers of some types of chemoreceptors.

In assembling the genome of *S. hermaphroditum*, we unexpectedly also were able to assemble genomes not only for its bacterial symbiont *X. griffiniae*, but also for eight bacterial genera that have been recurrently found associated with *Steinernema* or *Heterorhabditis* (Figure 8). The concurrence of these genera with past observations suggests that these microbes may be playing a biologically important role that underscores their consistently observed association with EPNs. However, the origin of these microbes and how they became associated with our laboratory strain of *S. hermaphroditum* PS9179 remains unknown. These nematodes were initially obtained from India, were subsequently grown on waxworms from various commercial sources, and were later grown from bleach-treated eggs on bacterial lawns of *X. griffiniae* prior to isolation of genomic DNA. It is unclear at what stage of this cultivation history or from what environmental source these associated microbes were acquired by *S. hermaphroditum*. However, our data, combined with previous reports of diverse types of microbes associated with EPNs, provide further impetus to explore the the microbiome composition, localization, and potential function of other laboratory strains of *Steinernema* or *Heterorhabditis* nematodes. Our genomic analysis of *S. hermaphroditum* and its associated bacteria provides a strong foundation for their use as a new biological model, and offers new avenues of exploration.

## Data availability

Genomic data (raw genomic and RNA-seq reads, genome assembly, and protein-coding gene predictions) for *S. hermaphroditum* have been archived in GenBank (BioProject PRJNA982879) and in WormBase ParaSite (https://parasite.wormbase.org/Steinernema_hermaphroditum_prjna982879/Info/Index). The *S. hermaphroditum* genome assembly has been archived in GenBank under the accession number GCA_030435675.1; *S. hermaphroditum* genomic sequencing reads have been archived at the SRA under accession numbers SRX20671584, SRX20671585, and SRX20671586; and *S. hermaphroditum* RNA-seq reads have been archived at the SRA under accession numbers SRX20672708 and SRX20672840. Supplementary Files S1 through S7 have been archived at the Open Science Framework under https://osf.io/3t5sc. Genomic data for microbial genomes and contigs will be archived in GenBank, MAGE, and *osf.io*.

## Methods

### Sample procurement, preparation and storage

Growth of *S. hermaphroditum* PS9179 was done as in Cao et al. (Cao et al. 2022).

### Genomic DNA extraction

To minimize heterozygosity in the genome assembly, the nematodes were inbred by selfing single hermaphrodites for at least 10 generations before samples were prepared for DNA extraction. The inbred infective juveniles (IJs) were cleaned using M9 buffer (composed of 6 g Na₂HPO₄, 3 g KH₂PO₄, and 5 g NaCl dissolved in 900 ml of distilled deionized water and autoclaved) and then transferred to a lawn of their symbiotic bacteria, *Xenorhabdus griffinae*, on nematode growth media (NGM) plates. The NGM plates were prepared using 20 g agar, 3 g NaCl, 10 ml uracil solution (2 g/L), 2.5 g peptone, and 975 ml of DI water, and autoclaved at 121°C under 15 psi pressure for 15 minutes. After autoclaving, 1 ml of cholesterol solution (5 mg/ml in ethanol), 1 ml of 1 M MgSO₄, 1 ml of 1 M CaCl₂, and 25 ml of 1 M KPO₄ buffer (pH 6.0) were added, and the plates were incubated at 27°C. The nematodes were cultured on NGM plates for 4-5 days until they contained a diverse mixture of life stages. For DNA extraction, the nematodes were surface sterilized by treating them with 15 ml of 4 mM Hyamine 1622 solution (Sigma, 51126) along with a bactericidal antibiotic cocktail consisting of carbenicillin (Fisher Scientific, BP2648), streptomycin (Fisher Scientific, BP910), and kanamycin, each at a concentration of 150 mg/ml, for 30 minutes. They were then washed 10 times with M9 solution and subjected to DNA extraction using the phenol-chloroform method.

### Genomic DNA sequencing

Genomic DNA was harvested largely by previous methods (Dillman et al. 2015) but with surface sterilization done by washing rather than bleaching. The choice of washing over bleaching was probably crucial for our being able to sequence and assemble associated bacterial genomes. Nanopore sequencing of genomic DNA was done at UC Riverside on a MinION with a FLO-MIN106D flow cell and with the kit type SQK-LSK110. Nanopore reads were decoded from native fast5 format into FASTQ format with guppy_basecaller from Guppy 5.0.11 (Oxford Nanopore Technologies; GPU-enabled linux64) using the arguments ‘*-r --num_callers 6 --cpu_threads_per_caller 1 --trim_barcodes --device auto*’. To lower the computational burden of further adapter-trimming, Nanopore reads were first size-selected with SeqKit2 2.0.0 (Shen et al. 2024) to ≥2.5 kb using the argument ‘*--min-len 2500*’; size-selected reads were trimmed for edge adapters and censored for internal adapters by porechop 0.2.4 (https://github.com/rrwick/Porechop), using the argument *’--discard_middle*’; they were then re-size-selected with SeqKit2 to ≥2.5 kb, yielding a final genome coverage of 137x. Illumina paired-end PCR-free shotgun of genomic DNA was done by Novogene; Illumina genomic reads were quality-trimmed, adapter-trimmed, and size-selected with fastp 0.20.1 (Chen et al. 2018), using the arguments ‘*--length_required 75*’, with a final genome coverage of 106x. Omni-C (non-restriction enzyme-dependent Hi-C) was done by Dovetail (now Cantata Bio); Omni-C (Illumina) reads were quality-trimmed, adapter-trimmed, and size-selected with fastp 0.23.2 (Chen et al. 2018), using the arguments ‘*--length_required 75 --max_len1 150 --detect_adapter_for_pe --dedup*’, with a final genome coverage of 392x.

### RNA harvesting and sequencing

For RNA collection, *S. hermaphroditum* PS9179 animals were grown on NGM agar media in Petri plates with lawns of *X. griffiniae* HGB2511 as a food source. Two samples were collected: a mixed-stage population, and dispersal-stage IJ larvae that were recovered in water traps adjacent to crowded populations of *S. hermaphroditum* nematodes on NGM agar media that had exhausted their lawns of HGB2511 bacteria. Total RNA was recovered by washing the collected nematodes several times in cold M9 buffer before resuspending the pelleted worms in Trizol. The Trizol suspension was frozen in a mortar using liquid nitrogen and ground into a fine powder using a pestle, maintained in its frozen state using repeated applications of liquid nitrogen. This powder was allowed to thaw, mixed with chloroform, and centrifuged; total RNA was recovered from the aqueous fraction by precipitation using isopropanol and ethanol. RNA-seq libraries were built using Illumina’s TruSeq RNA Sample Prep Kit v2 executed according to manufacturer’s instructions, using 1 µg of total RNA for each sample. RNA-seq libraries were sequenced in paired-end mode with read lengths of 150 nt. Newly generated *S. hermaphroditum* RNA-seq libraries are listed in Supplementary Table S1.

### General computation

When possible, we used bioconda (https://bioconda.github.io) to install and run version-controlled instances of software (Gruning et al. 2018). For reformatting or parsing of computational results, we used Perl scripts either developed for general use or custom-coded for a given analysis. All such Perl scripts (named below with italics and the suffix “*.pl*”) were archived on GitHub (https://github.com/schwarzem/ems_perl). Sources of software programs used in this study are listed in Supplementary Table S6.

### Nematode genomes, transcriptomes, coding sequences, and proteomes

For transcriptomic or proteomic analyses, published genome sequences, coding sequences, proteomes, and gene annotations of relevant nematodes were downloaded from WormBase (Sternberg et al. 2024) or WormBase ParaSite (Howe et al. 2017); they are listed in Supplementary Table S7. To extract protein and coding DNA (CDS DNA) sequences for protein-coding genes from a GTF or GFF3 file, we used gffread 0.12.7 (Pertea and Pertea 2020) with the arguments ‘*-g [input genome sequence FASTA file] -o /dev/null -C --keep-genes -L -P -V -H -l 93 -x [output CDS DNA FASTA file] -y [output protein FASTA file] [input GTF/GFF3 file]*’.

### Initial RNA-seq and cDNA assembly

Before using them for gene prediction or gene expression analysis, we trimmed adapters and low-quality sequences from Illumina RNA-seq reads with fastp 0.23.1 (Chen et al. 2018) using the arguments ‘*--length_required 75 --max_len1 150 --detect_adapter_for_pe*’. Before using them for cDNA assembly, fastp was run as above but with the added argument ‘*--dedup*’; we then assembled these RNA-seq data into cDNA sequences with Trinity 2.12.0 (Grabherr et al. 2011) using default arguments.

### Genome assembly

We assembled adaptor-trimmed and size-selected Nanopore reads with Raven 1.7.0 (Vaser and Šikić 2021) using default arguments. Among the resulting contigs, we identified contigs that contained microbial rather than nematode DNA with sourmash (2020-long-read-assembly-decontam version) (Pierce et al. 2019). Since microbial contigs in our assembly were not mere contaminants, but could represent symbiotic or commensal bacterial species of interest, we kept both the non-microbial (nematode) and the microbial contigs and error-corrected them in parallel. Because there is no uniformly agreed-upon recipe for error-correcting third-generation genome assemblies, we assayed the quality of the initially decontaminated *S. hermaphroditum* assembly, its successive correction products, and its final version with BUSCO 5.2.2 (Waterhouse et al., 2018) using the arguments ‘*--lineage_dataset nematoda_odb10 --mode genome*’, as well as with error-frequency scores determined by POLCA from MaSuRCA 4.0.8 (Zimin and Salzberg 2020) in later steps of error-correction. By iteratively assaying quality after each correction step, and only keeping correction steps that improved assembly quality, we empirically determined an optimal correction protocol for the *S. hermaphroditum* assembly, which we also applied to microbial contigs. We first error-corrected Raven contigs with Nanopore reads of ≥2.5 kb via one round of Racon 1.4.20 (Vaser et al. 2017) using the arguments ‘*-m 8 -x -6 -g -8 -w 500*’, in conjunction with minimap2 2.24 (Li 2018) using the argument ‘*-ax map-ont*’. We then error-corrected contigs with Nanopore reads of ≥2.5 kb via one round of medaka_consensus from medaka 1.5.0 (*https://github.com/nanoporetech/medaka*) using the argument ‘*-m r941_min_hac_g507*’. We finally error-corrected contigs with paired-end fastp-filtered Illumina reads via one round of POLCA in conjunction with BWA 0.7.17 (Li 2013), both using default arguments. Microbial contigs were then analyzed without further scaffolding.

### Chromosomal identification

Having error-corrected the *S. hermaphroditum* contigs, we chromosomally scaffolded them with fastp-trimmed Omni-C reads: we first ran juicer.sh from Juicer 1.6 (Durand et al. 2016b) using the argument ‘*-s none*’; we then ran run-asm-pipeline.sh from 3D-DNA 180922 (Dudchenko et al. 2017) using the arguments ‘*--mode haploid --sort-output --input 1500 --editor-repeat-coverage 6*’, in conjunction with juicer_tools.jar from Juicebox 2.13.07 (Durand et al. 2016a), BWA 0.7.17, SAMtools 1.9 (Danecek et al. 2021), and LASTZ 1.04.15 (Harris 2007). We identified which chromosomal scaffolds corresponded to which nematode chromosomes both by syntenic comparisons to published chromosomal nematode genome assemblies with promer and mummerplot in MUMMER4 4.0.0rc1 (Marcais et al. 2018) using the respective arguments ‘*--mum --minmatch 30*’ and ‘*--filter --terminal png --size large*’, as well as by visualizing Nigon elements of syntenic genes conserved among nematode chromosomes (Gonzalez de la Rosa et al. 2021); visualizations were generated with vis_ALG (*https:// github.com/pgonzale60/vis_ALG* and https://pgonzale60.shinyapps.io/vis_alg). Scaffolds were accordingly renamed to achieve clear synteny with published nematode chromosomes.

### Repetitive DNA element prediction and annotation

We generated an initial set of repetitive DNA elements from our fully-error-corrected *S. hermaphroditum* contigs with RepeatModeler2 2.0.3 (Flynn et al. 2020) and the DNA repeat library DFAM 3.6 (Storer et al. 2021), in conjunction with the programs CD-HIT 4.8.1 (Fu et al. 2012), GenomeTools 1.5.9 (Gremme et al. 2013), LTR_retriever 2.9.0 (Ou and Jiang 2018), MAFFT 7.505 (Nakamura et al. 2018), NINJA 0.98 (Wheeler 2009), Python 3.9.7, RECON 1.08 (Bao and Eddy 2002), RepeatScout 1.0.6 (Price et al. 2005), RMBlast 2.11.0 (Tempel 2012), TRF 4.0.9 (Benson 1999), and UCSC TwoBit tools (version kent_v362). We first ran the BuildDatabase program of RepeatModeler2 with the argument ‘*-engine rmblast*’, and then ran its RepeatModeler program with the argument ‘*-LTRStruct*’. In practice, some elements identified by RepeatModeler could be tandemly repeated gene families (either protein-coding genes, or ncRNAs such as rDNA). To detect and remove such sequences, we identified putative repeat elements likely to encode either protein-coding or ncRNA-coding genes. To detect possible protein-coding genes, we translated putative repeat elements into peptides with getorf from EMBOSS 6.6.0 (Rice et al. 2000) using the argument ‘*-minsize 90*’. We used these peptides to search Pfam 35.0 (Mistry et al. 2021) with hmmscan from HMMER 3.3.2 (Eddy 2009) using the argument ‘*--cut_ga*’; we also used these peptides to search *C. elegans* proteins from WormBase release WS285 with BlastP from BLAST 2.10.1 (Camacho et al. 2009) using the arguments ‘*-evalue 1e-06 -outfmt 7 -seg yes*’. To detect possible ncRNA-coding genes, we directly scanned putative repeat elements with cmsearch from INFERNAL 1.1.4 (Nawrocki and Eddy 2013) and the ncRNA motif library Rfam 14.8 (Kalvari et al. 2021) using the argument ‘*--cut_ga*’. As a negative control, we ran equivalent HMMER, BLAST, and INFERNAL searches on known *C. elegans* transposon sequences from WormBase WS285 and known *C. elegans* repetitive DNA elements from DFAM 3.6 (RepeatMasker.lib); any protein or ncRNA matches in these *C. elegans* sequences could be ignored when also found in *S. hermaphroditum* sequences. Finally, we checked repetitive elements for annotations by RepeatModeler that clearly identified them as recognizable members of known repetitive element families (e.g., “DNA”, “LINE”, “LTR”, or “SINE”). Having identified elements that had HMMER, BLAST, and INFERNAL hits indicating that they were possible protein- or ncRNA-coding genes and that did not have unambiguous RepeatModeler annotations, we subtracted them to generate a final set of *S. hermaphroditum* repetitive DNA elements. We softmasked the genome assembly with its final set of repetitive DNA elements via RepeatMasker 4.1.2-p1 (https://www.repeatmasker.org) using the arguments ‘*-s -xsmall -gccalc -gff*’.

### Protein-coding gene prediction

To predict protein-coding genes in *S. hermaphroditum*, we ran *braker.pl* in BRAKER2 2.1.6 (Brůna et al. 2021) on our repeat-softmasked chromosomal *S. hermaphroditum*. BRAKER2 requires an input genome sequence to have its FASTA header lines previously stripped of comments; we did this with *uncomment_FASTA_headers.pl*. To guide BRAKER2 gene predictions, we used both transcriptomic and proteome sequence data. For transcriptomic data, we mapped *S. hermaphroditum* RNA-seq reads to the genome assembly with HISAT2 2.2.1 (Kim et al. 2019) and reformatted them into a sorted and indexed BAM alignment with SAMtools 1.14 (Danecek et al. 2021) using default arguments. For proteomic data, we downloaded protein sequences for *Steinernema* and other clade IV nematode species from WormBase ParaSite WBPS17 (Howe et al. 2017): *Bursaphelenchus okinawaensis* (Sun et al. 2020); *Panagrellus redivivus* (Srinivasan et al. 2013); *S. carpocapsae* (Serra et al. 2019); *S. feltiae* (Fu et al. 2020), *S. glaseri*, *S. monticolum*, and *S. scapterisci* (Dillman et al. 2015); *Rhabditophanes diutinus* KR3021 and *S. ratti* (Hunt et al. 2016). Running BRAKER2 also required us to install: AUGUSTUS 3.4.0 (Stanke et al. 2008); BamTools 2.5.2 (Barnett et al. 2011); cdbfasta 1.00 (Pertea et al. 2003); DIAMOND 0.9.24.125 (Buchfink et al. 2021); GeneMark-ES/ET 4.71 (Lomsadze et al. 2014); GenomeThreader 1.7.3 (Gremme et al. 2005); MakeHub 1.0.6 (Hoff 2019); ProtHint 2.6.0 (Bruna et al. 2020); and SAMtools 1.15.1 (Danecek et al. 2021).

We ran BRAKER2 four times in parallel with different input data sets and arguments, before merging the four predicted gene sets with TSEBRA 1.0.3 (Gabriel et al. 2021): the first BRAKER2 run used RNA-seq data alone, with default arguments; the second run used proteomic data alone, with the argument ‘*--epmod*’; the third run used RNA-seq, proteomic, and repeatmasking data, with the arguments ‘*--etpmode --softmasking*’; the fourth run used *ab initio* predictions, with the argument ‘*--esmode*’. After running BRAKER2, we merged predictions into a single gene set with TSEBRA using the argument ‘*-k*’ to keep all transcripts, and renamed gene, transcript, and exon names with a modified version of *updateBRAKERGff.py* (https://github.com/Gaius-Augustus/BRAKER/issues/416) and custom Perl one-line commands. To identify redundant transcripts that encoded identical proteins (either as complete or subset sequences) we used CD-HIT 4.8.1 with the arguments ‘*-g 1 -M 0 -d 100 -c 1.0*’ and censored all redundant transcripts encoded by a common gene, while retaining at least one transcript for each predicted gene.

### ncRNA locus identification

To identify ncRNA gene loci, we searched the *S. hermaphroditum* genome with cmscan from INFERNAL 1.1.5 (Nawrocki and Eddy 2013) and the ncRNA motif library Rfam 14.10 (Kalvari et al. 2021) using the arguments ‘*--rfam --cut_ga --nohmmonly --clanin Rfam-14.10.clanin --tblout [tabular output] --fmt 2 -o /dev/null*’. We converted the search results to GFF3 format with *infernal-tblout2gff.pl* (https://github.com/nawrockie/jiffy-infernal-hmmer-scripts) using the arguments ‘*--cmscan --fmt2 --all*’., and further reformatted them with *fix_infernal_gff_26apr2024.pl*. We further converted these into nonredundant genomic loci by the following general procedure.

### Mapping genome annotations to nonredundant genome regions

To map gene annotations (such as Rfam hits or repetitive elements) into nonredundant genomic regions, we converted their annotation file from GFF to BED file format with *gff2bed* from BEDOPS 2.4.41 (Neph et al. 2012) using the argument ‘*--do-not-sort*’, sorted the BED file with ‘*sort -k1,1 -k2,2n*’ in Linux, consolidated the BED file’s genomic intervals with the merge program of BEDTools 2.30.0 (Quinlan and Hall 2010) using the argument ‘*-delim “|”*’. Where a GFF annotation file was needed, the consolidated BED file was converted back to GFF format with convert_bed2gff.pl from AGAT 1.3.3 (Dainat 2024). Source and sequence type annotations in the resulting GFF were converted from ‘data\tgene’ to ‘maf-convert\tregion’ with inline Perl.

### Assessing protein-coding genes

We determined general statistics of protein-coding gene products with *count_fasta_residues.pl* using the arguments ‘*-b -e -t prot*’; we obtained gene counts from maximum-isoform proteome subsets generated with *get_largest_isoforms.pl* using the arguments ‘*-t parasite*’ or *’-t maker*’. We determined and compared the completeness of predicted *A. ceylanicum* protein-coding gene sets with BUSCO 5.2.2 using the arguments ‘*--lineage_dataset nematoda_odb10 --mode proteins*’ (Waterhouse et al. 2018).

### GenBank deposition

We used NCBI’s table2asn 1.27.792 (Sayers et al. 2024) to convert the final genome assembly and protein-coding gene set for uploading to GenBank (Sayers et al. 2024), with the arguments ‘*-M n -J -c b -euk -locus-tag-prefix QR680 -gaps-min 1 -l proximity-ligation -Z -V b*’. Before doing this, we reformatted gene predictions into standardized GFF3 format with *agat_convert_sp_gxf2gxf.pl* from AGAT 1.1.0 (Dainat 2024). We also identified transcripts with ambiguous translations, by comparing protein translations by gffread to those by table2asn; any transcript that could not be identically translated by both programs was censored, as was any gene with only ambiguous translations. We identified any remaining microbial contigs either through the NCBI Foreign Contamination Screen (Astashyn et al. 2024) or through OrthoFinder matches to bacterial genes; these contigs were either censored before submission to GenBank or suppressed after it. The final set of genes and transcripts was uploaded to GenBank, and used for subsequent analyses.

### Protein-coding gene annotation

For the protein products of our *S. hermaphroditum* gene set, we predicted both N-terminal signal sequences and transmembrane alpha-helical anchors with Phobius 1.01 (Käll et al. 2004). We predicted coiled-coil domains with Ncoils 2002.08.22 (Lupas 1996). We predicted low-complexity regions with PSEG 1999.06.10 (Wootton 1994). We identified protein motifs with *hmmscan* in HMMER 3.4 (Eddy 2009) and Pfam 36.0 (Mistry et al. 2021), using the argument ‘*--cut_ga*’ to impose family-specific significance thresholds and ‘*--tblout*’ to export tabular outputs; we also identified protein motifs with *interproscan.sh* in InterProScan 5.67-99.0 (Blum et al. 2021) using the arguments ‘*-dp -iprlookup -goterms*’. We predicted Gene Ontology (GO) terms (Carbon et al. 2021) with EnTAP 1.0.1 (Hart et al. 2020) using the argument ‘*--runP*’, and using selected UniProt (UniProt 2023) and RefSeq (O’Leary et al. 2016) proteome databases from highly GO-annotated model organisms (Supplementary Table S7); proteome databases were generated with *makedb* from DIAMOND 0.9.9 (Buchfink et al. 2021), itself bundled with EnTAP. For our comparisons of Pfam motif frequencies between nematodes, we identified Pfam 36.0 motifs in proteomes of four other nematodes (*C. elegans*, *Bursaphelenchus xylophilus*, *Heterorhabditis bacteriophora*, *S. carpocapsae*, and *S. ratti*) by the same method as in *S. hermaphroditum*. We identified orthologies between *S. hermaphroditum* and other nematodes with OrthoFinder 2.5.5 (Emms and Kelly 2019) using the arguments ‘*-S diamond_ultra_sens - M msa*’. to generate both orthology tables and species phylogenies, or ‘*-S diamond_ultra_sens-og*’ to generate orthology tables alone. Because the granularity of gene orthology groups produced by OrthoFinder varies with the number of species analyzed, we ran OrthoFinder analyses with varying numbers of input species.

### Statistical significance of differing protein motif-encoding genes

To identify significant differences between sets of genes encoding Pfam 36.0 protein motifs in *S. hermaphroditum* versus other nematodes, we first reformatted Pfam results for each nematode into a gene-centric tab-delimited format with *pfam_hmmscan2gene_freqs.pl*, and merged species pairs of the reformatted results with *merge_pfam2gene_freqs.pl*. We then used *motif_group_fisher_12sep2024.pl*, which computed p-values from two-tailed Fisher tests with the Perl module *Text::NSP::Measures::2D::Fisher::twotailed*; for multiple hypothesis testing (e.g., comparisons to sets of protein motifs) this Perl script also computed q-values via the *qvalue* program of the MEME 5.4.1 software suite (Bailey et al. 2009).

### Phylogenetic trees and time estimates

As part of our OrthoFinder analyses, species trees were automatically computed with STAG (Emms and Kelly 2018) and rooted with STRIDE (Emms and Kelly 2017); a 16-taxon tree generated in this way was used for CAFE analysis.

To estimate phylogenetic divergence times for species within the nematode phylum, we instead began with the multiple sequence alignment file (MSA) generated by a 22-species OrthoFinder analysis (SpeciesTreeAlignment.fa). We used this MSA to compute a species phylogeny with IQ-TREE 2.2.0.3 (Minh et al. 2020), using the arguments ‘*-s SpeciesTreeAlignment.fa -m MFP -nt AUTO -ntmax 64 -bb 1000 -alrt 1000*’, and using the resulting consensus tree file (SpeciesTreeAlignment.fa.contree) for divergence time estimation. We extracted reference times for nematode species divergence from https://timetree.org (Kumar et al. 2022), which gave confidence intervals (CIs) for divergence times of two species pairs: between *C. elegans* and *C. briggsae*, we obtained a CI of 18.6 to 101.5 Mya, derived from (Blair et al. 2005; Cutter 2008); and between *C. elegans* and *P. pacificus*, we obtained a CI of 180.7 to 191.0 Mya (Douzery et al. 2004; Rota-Stabelli et al. 2013). To specify the model of amino acid substitution for estimating phylogeny, we downloaded mtREV24.dat from https://github.com/abacus-gene/paml/tree/master/dat. We then used mcmctree and codeml from PAML 4.9 to estimate species divergence times through Bayesian MCMC (Yang and Rannala 2006; Rannala and Yang 2007), with details given in Supplementary File S1.

### Visualizing chromosomal contents

We linearly plotted nematode chromosomes and their contents with JBrowse2 2.18.0 (Diesh et al. 2023). To visualize individual genes, we used GFF3 annotation files. To visualize chromosomal densities of genes or repetitive DNA regions, we used BigWig annotation files produced as follows: we used the genomecov program of BEDTools 2.30.0 (Quinlan and Hall 2010) to convert a BED annotation file to BEDGRAPH format; we sorted the BEDGRAPH file with ‘*sort -k1,1 -k2,2n*’ in Linux; and we converted the sorted BEDGRAPH file into BigWig format with bedGraphToBigWig from Kent software.

### Visualizing orthology memberships

We drew patterns of multispecies orthology group membership with UpSetR (Conway et al. 2017) via its Shiny app (https://gehlenborglab.shinyapps.io/upsetr), after extracting subsets of our 22-species OrthoFinder groups and reformatting them into a binary table with *pre.upset.orths_23dec2024.pl*.

### Visualizing phylogenies

We drew phylogenies with FigTree 1.4.4 (http://tree.bio.ed.ac.uk/software/figtree). For our phylogeny giving estimated divergence times, we displayed node ages under the Node Label parameter and added a scale bar, using the reverse axis setting on the scale axis; for the node bars, we applied a 95% confidence interval; and for the time scale bar, we set the scale factor to −1.

### Identifying evolutionarily expanding or contracting protein-coding gene families

Starting from a 16-taxon OrthoFinder analysis, we converted its species tree into an ultrametric tree with ape 5.8 (Paradis and Schliep 2019), and converted its orthology group table into a table of gene counts for each orthology group and gene/species combination with data.table 1.15.4. These file conversions were generated with an R script, provided as Supplementary File S2. We then used CAFE 5.1.0 (Mendes et al. 2021) with the arguments ‘*-k 3 -z*’) to identify gene families (orthology groups) from an OrthoFinder analysis whose gene number expanded or contracted over the ultrametric species tree derived from that OrthoFinder analysis. We visualized the results and summarized family size changes with CafePlotter (https://github.com/moshi4/CafePlotter), using CafePlotter v0.2.0 to generate all outputs except a visual summary of gene families with statistically significant changes (summary_significant_gene_family) provided by CafePlotter v0.1.1. This visual summary gives all families with any p-value up to 1; we generated it as an SVG file and manually corrected it with Adobe Illustrator, after scanning the CafePlotter output ‘result_summary.tsv’ with *filt_sig_cafe_02jan2025.pl* to only count families with p ≤ 0.05 for each phylogenetic node. We likewise used *filt_sig_cafe_02jan2025.pl* to identify specific gene families expanding or contracting at the *Steinernema* root node or *S. hermaphroditum* stem node.

### RNA-seq expression values

We quantitated gene expression from RNA-seq data sets with Salmon 1.10.2, generating expression values in transcripts per million (TPM) and estimating mapped read counts per gene (Patro et al. 2017). To prevent spurious mappings of RNA-seq reads, we used full selective alignment to a gentrome (a CDS DNA set, treated as a target for real mappings, combined with its genome, treated as a decoy for spurious mappings), followed by quantification using Salmon in non-alignment mode (Srivastava et al. 2020; Srivastava et al. 2021). For Salmon’s index program, we used the arguments ‘*--keepDuplicates -t [gentrome sequence] -d [decoy list]*’; for Salmon’s quant program, we used the arguments ‘*--seqBias --gcBias --posBias --libType A --geneMap [transcript-to-gene table] --mates1 [R1 reads] --mates2 [R2 reads]*’. Results from quant.genes.sf output files were reformatted with *make_salmon_TPM_slices.pl*.

### Identifying venom ortholog genes and their genomic clusters

We extracted lists of venom genes for *S. carpocapsae* from Lu et al. (Lu et al. 2017) and for *S. feltiae* from Chang et al. (Chang et al. 2019). Starting from our 4-species OrthoFinder analysis, we used *mark_venom_orths_11dec2024.pl* and these gene lists to annotate orthology groups with venom genes for each *Steinernema* species. From these venom-annotated orthology groups, we identified 567 *S. hermaphroditum* genes with orthology to a venom gene from either *S. carpocapsae* or *S. feltiae*, and 307 genes whose orthologs in either *S. carpocapsae* or *S. feltiae* consisted exclusively of venom genes. Of these two ortholog sets, 565 and 305 genes (respectively) were on chromosomal scaffolds and thus could be analyzed for genomic clustering, while two genes were on the non-chromosomally-assigned scaffold *Sherm_2022.07.24.06*. We used *gff_gene_subset_09nov2024a.pl* to extract GFF3 files with gene coordinates for the 565 or 305 *S. hermaphroditum* venom gene orthologs from our full gene-prediction GFF3. We then used GALEON V1 (Pisarenco et al. 2024) to determine whether these sets of 565 or 305 venom gene orthologs formed clusters in the *S. hermaphroditum* genome. We ran GALEON_ControlScript.py with the arguments ‘*clusterfinder -a GFFs/ -e disabled -g 50 -o [venom_set]*’; we ran GALEON_SummaryFiles.py with the arguments ‘*-fam [venom_set] -clust clusterfinder_Results_Directory/ -coords GFFs -ssize [chromosomal sizes table] -sfilter 7*’; and we ran GALEON_Report.py with the arguments ‘*-clust clusterfinder_Results_Directory/ -ssize [chromosomal sizes table] -echo False*’.

### Identifying possible microbial Nanopore reads

To identify which Nanopore reads were likely to come from microbes rather than *S. hermaphroditum*, we first mapped them with minimap2 2.24 to our microbial contigs, using the arguments ‘*-a -x map-pb*’. We extracted and pooled reads that either mapped to our microbial contigs or failed to map to them. Reads that did not map to our microbial contigs were then mapped as above to our *S. hermaphroditum* assembly, and any that failed to map were extracted and pooled with our microbially mapping reads to yield a single Nanopore read set. We used SAMtools 1.17 to identify and extract reads into a BAM alignment file by running ‘*samtools view -F 4 -b*’ (to select only mapping reads) or ‘*samtools view -f 4 -b*’ (to select only nonmapping reads); we then ran ‘*samtools fastq*’ to extract reads in FASTQ format from a given BAM.

### Identifying S. hermaphroditum-associated microbial sequences

We taxonomically identified both assembled microbial contigs and raw sequencing reads that could map to those contigs. We split a FASTA file of error-corrected non-nematode contigs into individual FASTA files with faSplit (version 377) on the Galaxy platform (Galaxy 2024). To determine the completeness of microbial contigs, we ran CheckM 1.0.18 (Parks et al. 2015); to taxonomically classify microbial contigs, we ran GTDB-Tk 1.7.0 (Chaumeil et al. 2019) using the GTDB database (Parks et al. 2022) and the “full tree” setting on the Kbase platform (Arkin et al. 2018). To taxonomically classify raw reads, we ran Kaiju (Menzel et al. 2016) on Kbase. (Norman et al. 2020). When we examined the taxonomic breakdown of the raw reads which failed to map to the *S. hermaphroditum* genome assembly, around 80% of the Illumina short reads were classified as *Xenorhabdus*; however, only about 30% of the Nanopore long reads were classified as *Xenorhabdus*, with a much larger fraction of the long reads classified as the other bacterial genera represented in our analysis.

## Acknowledgments

This work was supported by NSF EDGE FGT 2128266 to H.G.-B., P.W.S., and A.R.D. We thank NSF ACCESS for a research allocation on the Pittsburgh Supercomputing Center Bridges-2 Regular Memory cluster (TG-MCB190010), Titus Brown for use of the UC Davis Farm computing cluster, and the Millard and Muriel Jacobs Genetics and Genomics Laboratory at the California Institute of Technology for sequencing and computational support.

---

**Supplementary Figure S1:**
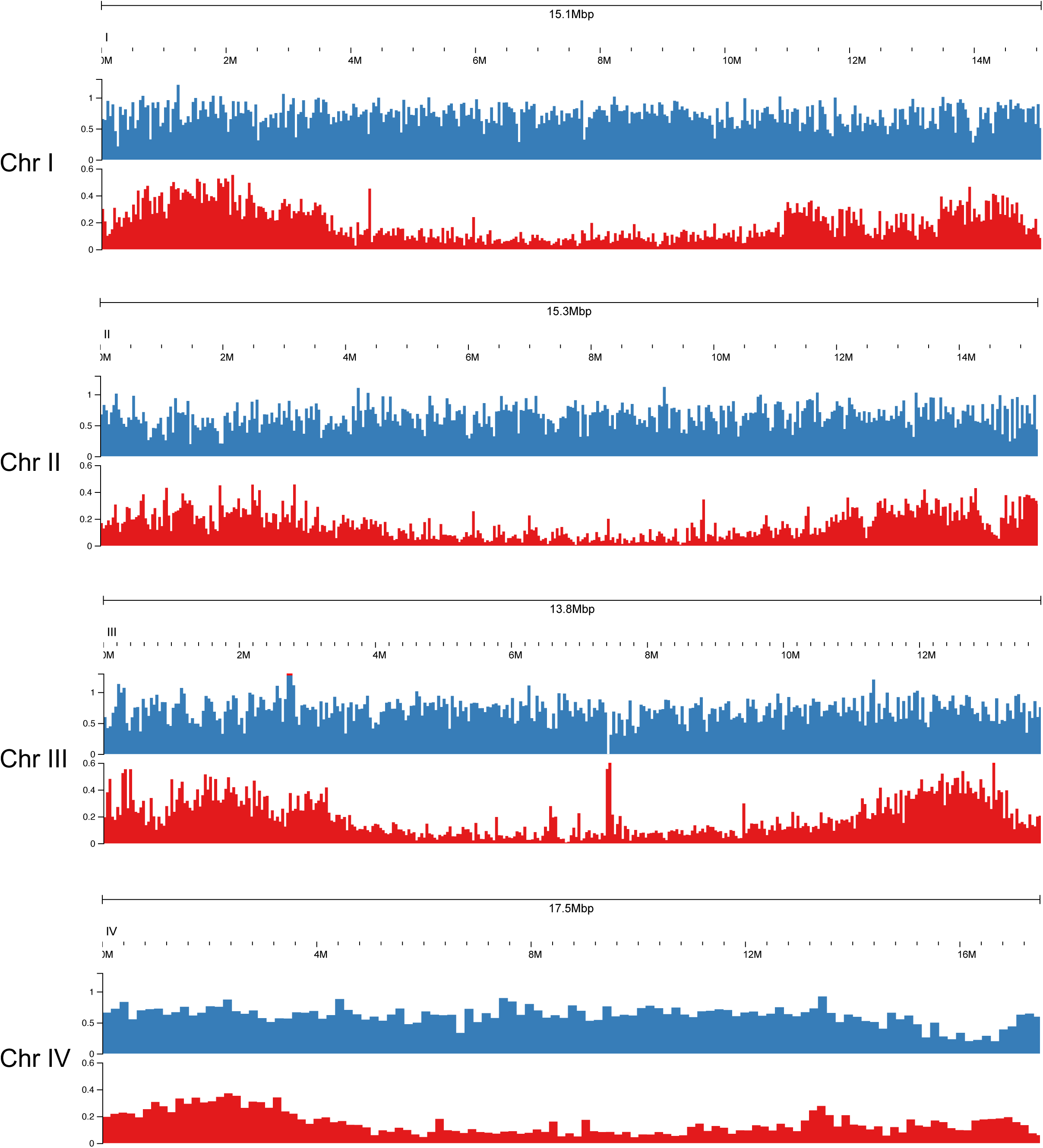
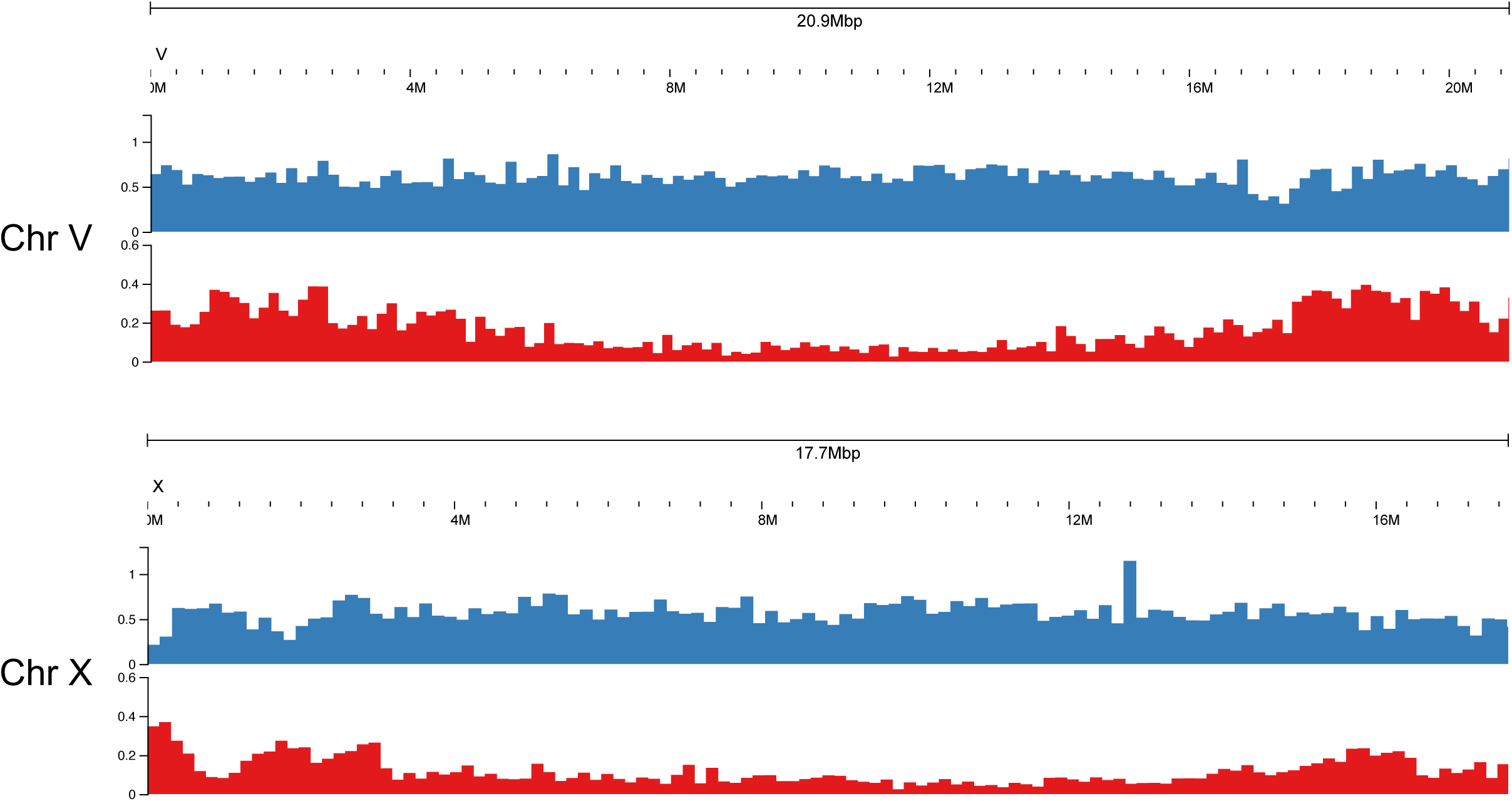
*C. elegans* chromosomal densities of genes and repetitive DNA. Densities of protein-coding genes (blue) and repetitive DNA regions (red). These were computed from existing *C. elegans* data (Supplementary Table S7) by the same methods as in Figure 3. Protein-coding prediction methods between *S. hermaphroditum* and *C. elegans* can be considered equivalent. However, the repetitive elements in *C. elegans* were defined by a very different process than we could use for *S. hermaphroditum*, and this should be kept in mind when comparing their chromosomal distributions.

**Supplementary Figure S2.**
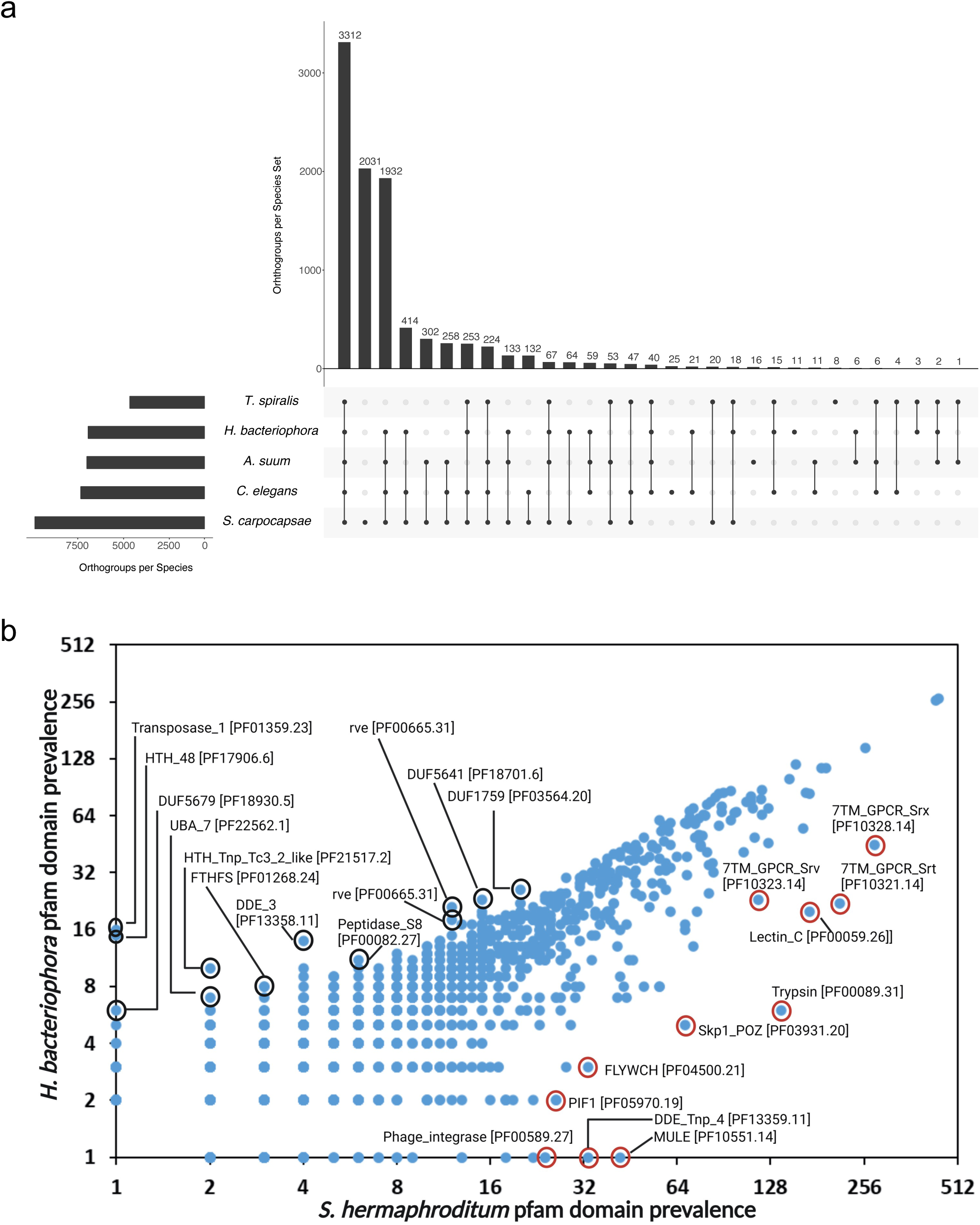
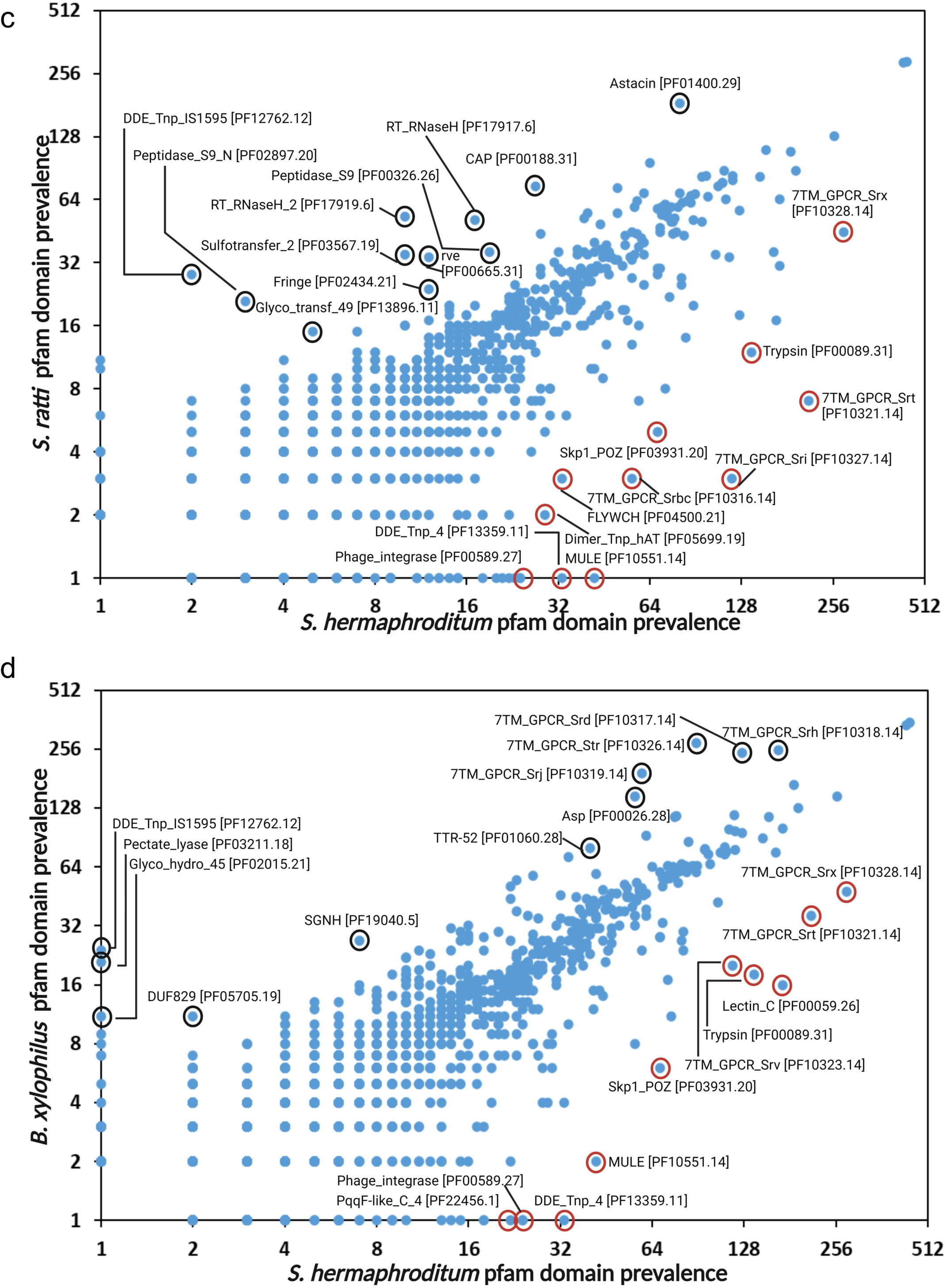
Orthologies and Pfam motif frequencies. (a) An UpSet plot (Conway et al. 2017) of overlapping orthologies between genes in *S. hermaphroditum* and five other representative nematode species (Figure 1b): *Trichinella spiralis* (clade I), *Ascaris suum* (clade III), *H. bacteriophora* (clade V), *C. elegans* (clade V), and *S. carpocapsae* (clade IV). Most orthology groups either involve all six species (3,312 orthologies), or only the close relative *S. carpocapsae* (2,031 orthologies), or all except the phylogenetically remote *T. spiralis* (1,932 orthologies). (b) Comparison of the frequencies of genes encoding Pfam motifs between *S. hermaphroditum* and *H. bacteriophora*. (c) Comparison of the frequencies of genes encoding Pfam motifs between *S. hermaphroditum* and *S. ratti*. (d) Comparison of the frequencies of genes encoding Pfam motifs between *S. hermaphroditum* and *B. xylophilus*. Selected motifs are circled, and are labeled with their Pfam abbreviations and accession numbers.

**Supplementary Figure S3:**
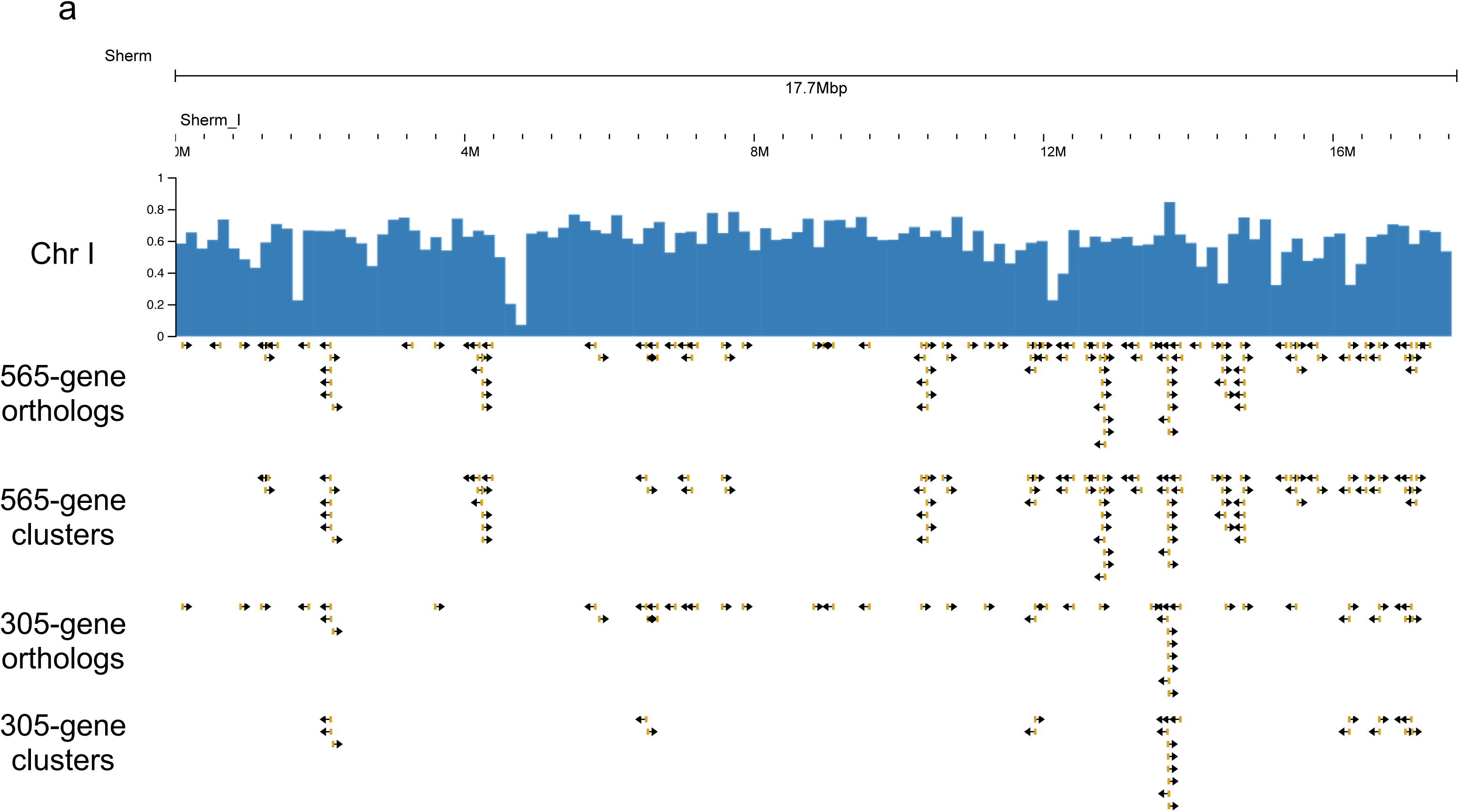
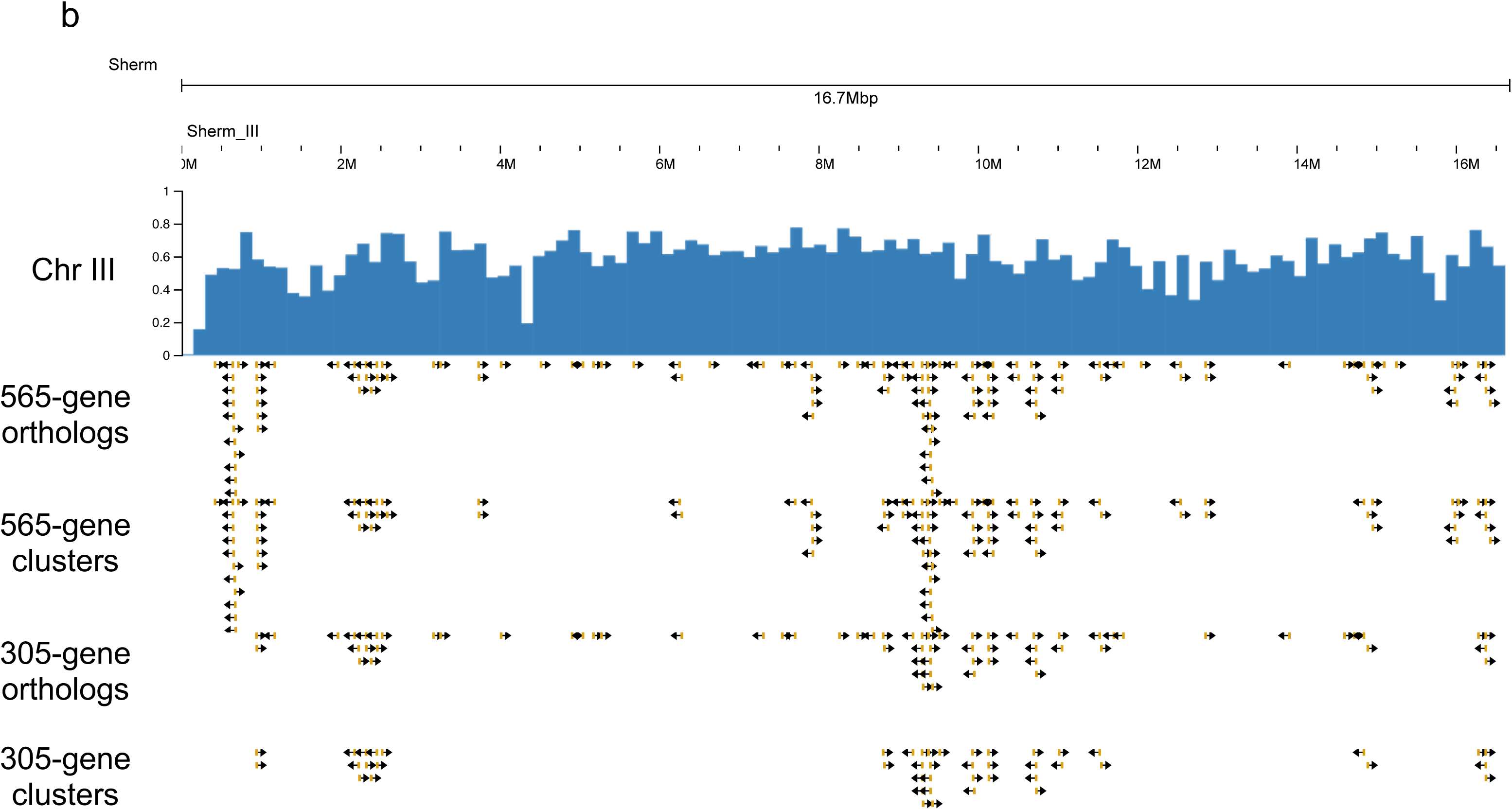
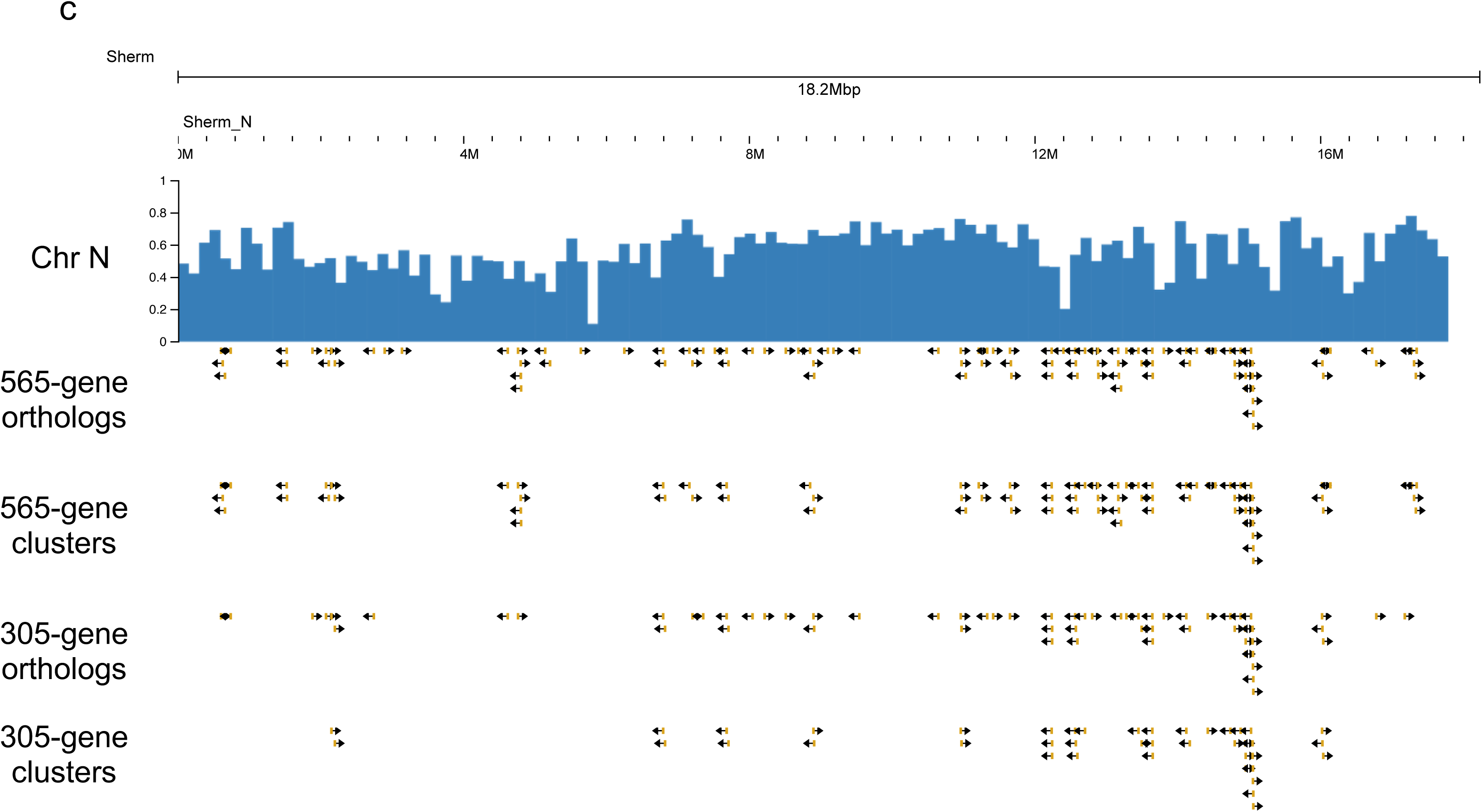
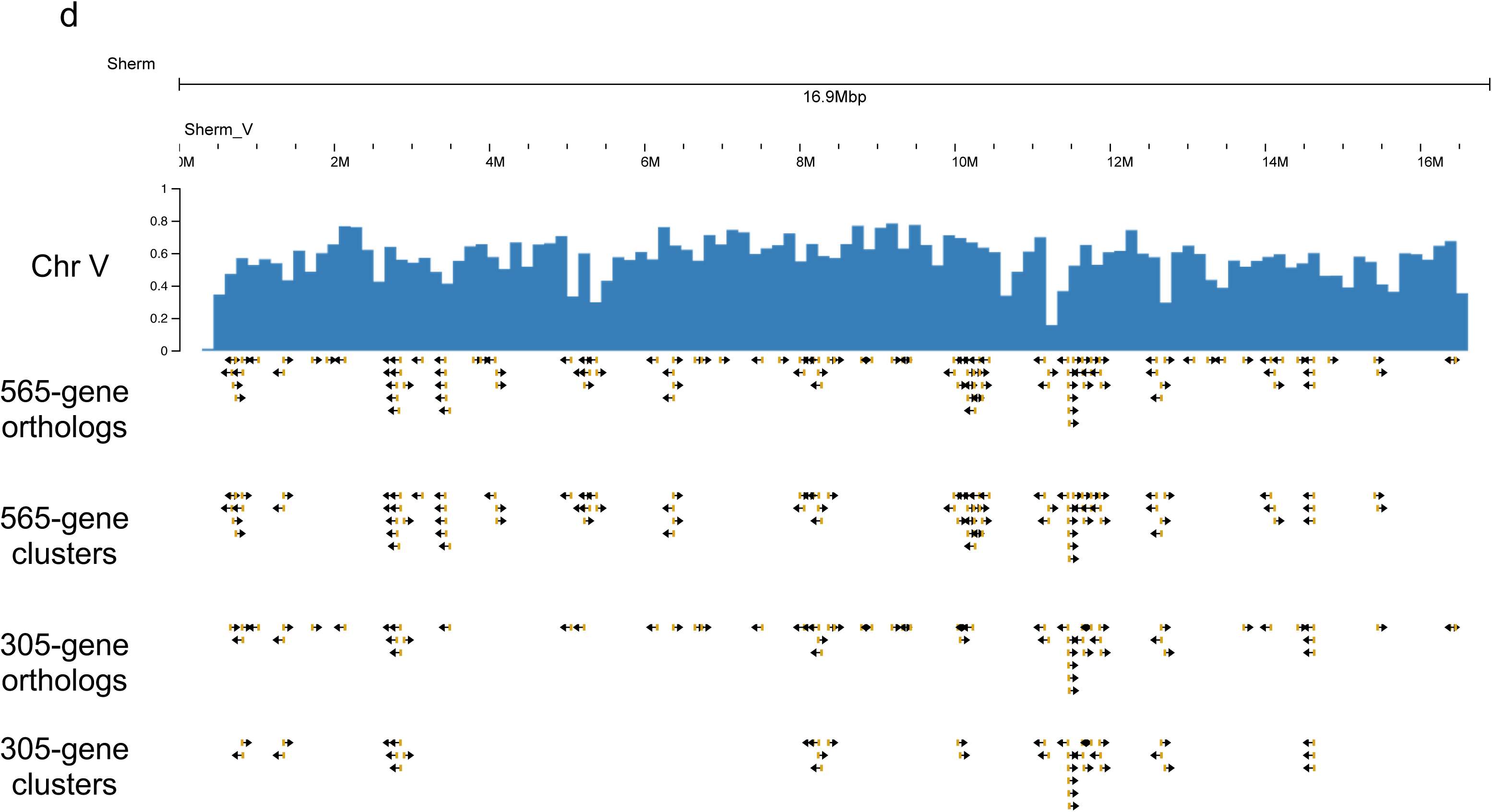
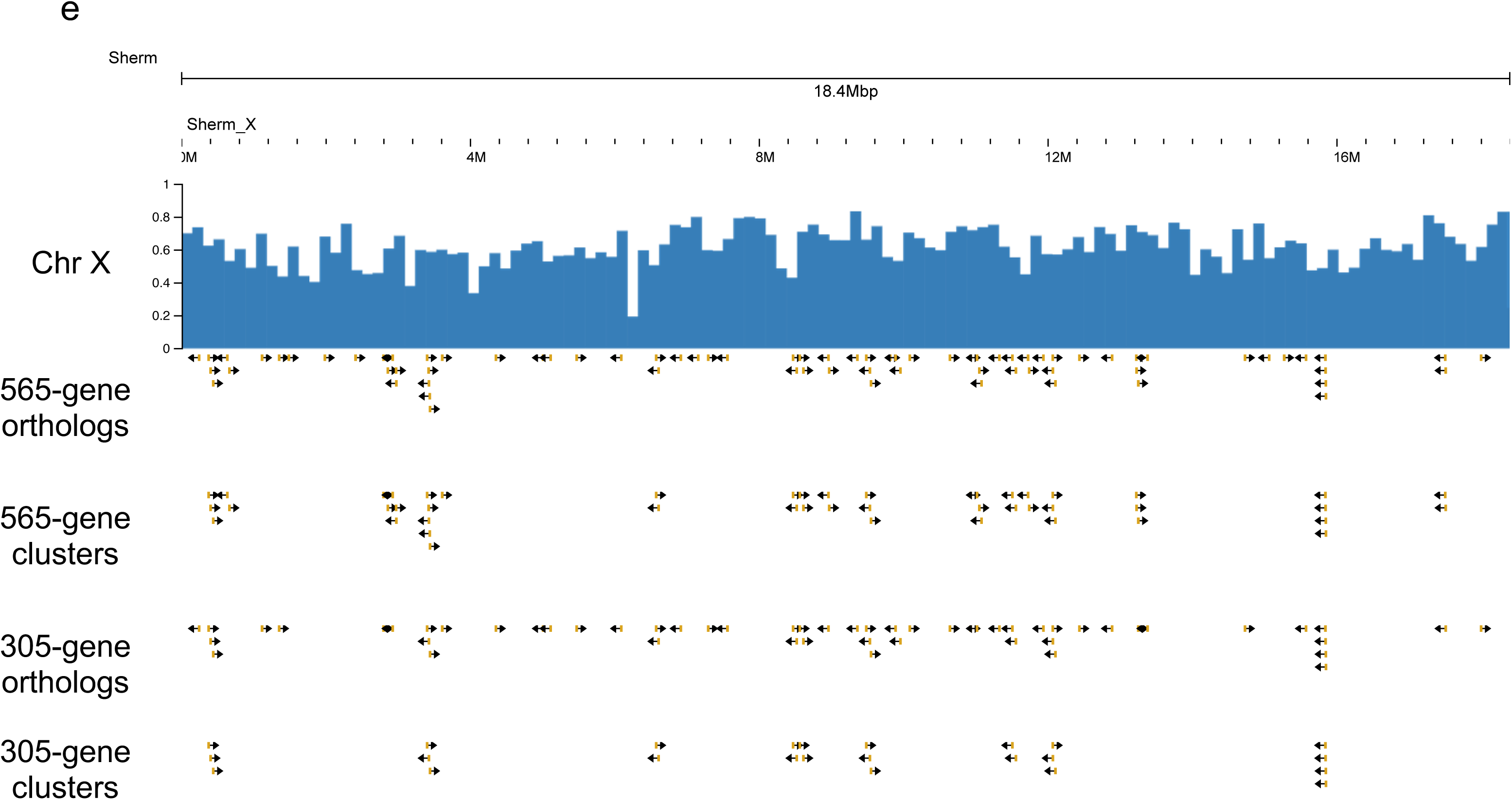
Venom ortholog genes and their clusters, from both the 565-gene and 305-gene venom ortholog sets. For each chromosome, the following are shown: mean gene density; a broadly defined set of 565 venom ortholog genes; clusters identified by GALEON (Pisarenco et al. 2024) from that 565-gene set; a narrowly defined set of 305 venom ortholog genes; and clusters identified by GALEON from that 305-gene set. (a) Chromosome I; (b) Chromosome III; (c) Chromosome N; (d) Chromosome V; and (e) Chromosome X.

**Supplementary Figure S4:**
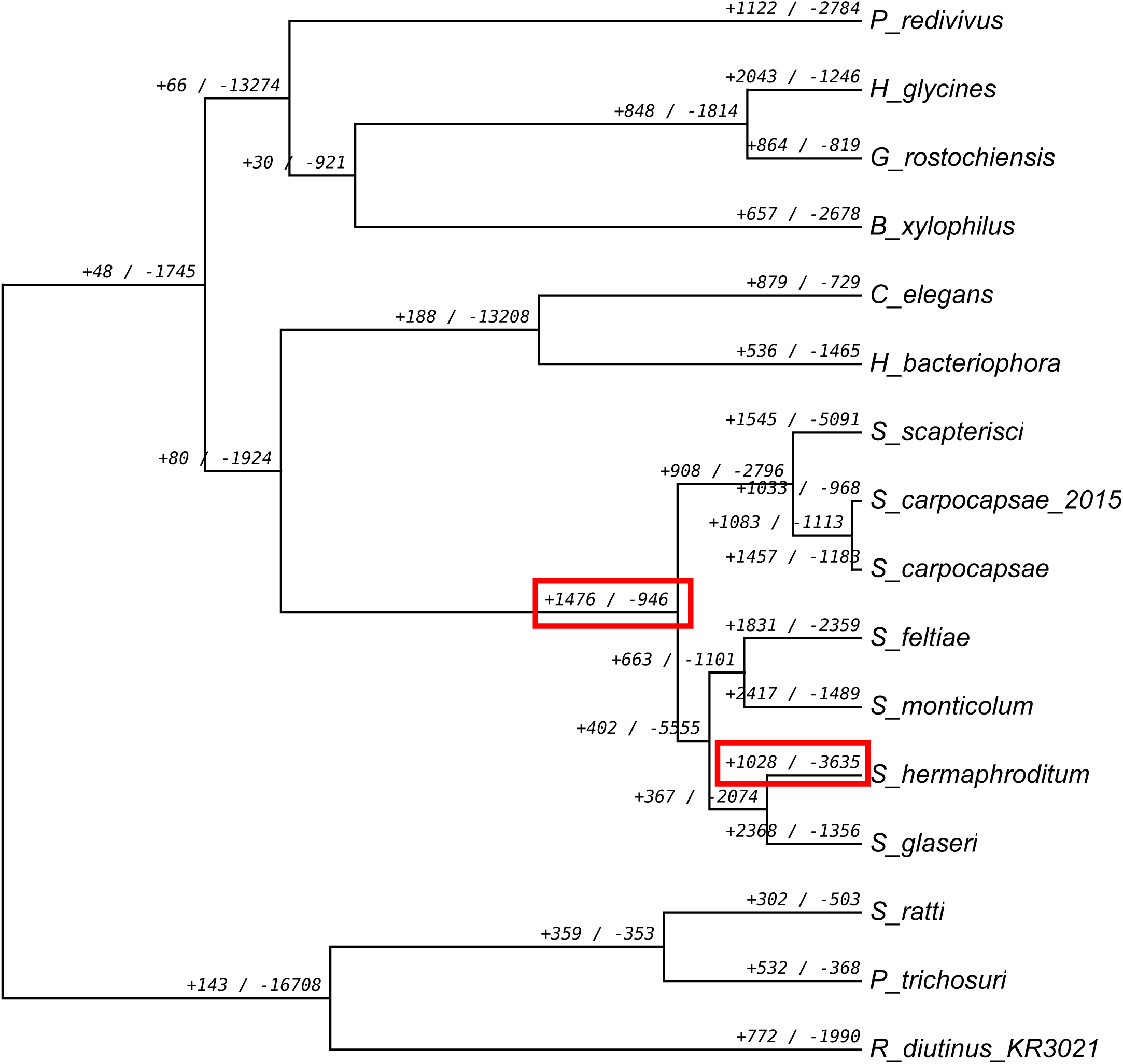
All gene families changing size in *Steinernema* or *S. hermaphroditum*. This shows all gene families that gene families that gained or lost genes at nodes of a phylogenetic tree of 16 nematode protein-coding gene sets, regardless of whether these changes were scored as statistically significant. It is otherwise equivalent to Figure 7.

**Supplementary Table S1: Genomic DNA and RNA-seq sequencing libraries generated in this study.** “Library ID” provides a short abbreviation for each library; “Description” summarizes biological contents. For each library, “SRA accession”, “BioProject accession” and “BioSample accession” provide accession numbers; “Number of reads”, “Total length in nt”, and “Mean read length in nt” give the number of reads, total sequence in nt, and mean read length.

**Supplementary Table S2: Protein-coding gene annotations for *Steinernema hermaphroditum*.** Its data columns are as follows.

**Gene:** Name of a predicted protein-coding gene in the *S. hermaphroditum* genome assembly. All further data columns are pertinent to that particular gene.

**UniProt_ID|GenBank_ID:** Identification numbers extracted from UniProt (UniProt 2023) and GenBank (Sayers et al. 2024).

**Strict_elegans_ortholog:** Strict (one-gene-to-one-gene) orthologies between a given *S. hermaphroditum* gene and its orthologous *C. elegans* gene; these were extracted from a 14-species OrthoFinder analysis (further annotated in **OFind_Summary_14spp** and **OFind_Full_14spp**).

**Max_prot_size:** The size of the largest predicted protein product.

**Prot_size:** This shows the full range of sizes for all protein products from a gene’s predicted isoforms. Phobius: This denotes predictions of signal and transmembrane sequences made with Phobius (Käll et al. 2004). ‘SigP’ indicates a predicted signal sequence, and ‘TM’ indicates one or more transmembrane-spanning helices, with N helices indicated with ‘(Nx)’. Varying predictions from different isoforms are listed.

**NCoils:** This shows coiled-coil domains, predicted by Ncoils (Lupas 1996). Both the proportion of such sequence (ranging from 0.01 to 1.00) and the exact ratio of coiled residues to total residues are given. Proteins with no predicted coiled residues are blank.

**Psegs:** This shows what fraction of a protein is low-complexity sequence, as detected by PSEG (Wootton 1994). As with Ncoils, relative and absolute fractions of low-complexity residues are shown.

**Pfam:** Predicted protein motifs from Pfam 36.0 (Mistry et al. 2021), with family-specific significance thresholds.

**InterPro:** Predicted protein motifs from InterProScan 5.67-99.0 (Blum et al. 2021).

**[X].TPM:** For each individual RNA-seq data set (with ‘X’ denoting the data set’s abbreviation), this gives gene expression levels in TPM, computed by Salmon (Patro et al. 2017). Here and for **[X].reads**, the abbreviations ‘Sherm_mixed_stage’ or ‘Sherm_IJ’ denote RNA-seq data from mixed-stage or infectious juvenile *S. hermaphroditum*.

**[X].reads:** For each individual RNA-seq data set (with ‘X’ denoting the data set’s abbreviation), this gives numbers of mapped RNA-seq reads per gene, computed for individual RNA-seq data sets by Salmon (Patro et al. 2017), with fractional values rounded down to integers.

**EggNOG_description:** EggNOG descriptions (Hernandez-Plaza et al. 2023), generated with EnTAP (Hart et al. 2020).

**EggNOG_KEGG:** KEGG codes (Kanehisa et al. 2023), generated with EnTAP (Hart et al. 2020). **GO_Biological:** Annotations from the biological subset of Gene Ontology (GO) terms (Carbon et al. 2021), generated with EnTAP (Hart et al. 2020).

**GO_Molecular:** Annotations from the molecular subset of Gene Ontology (GO) terms (Carbon et al. 2021), generated with EnTAP (Hart et al. 2020).

**GO_Cellular:** Annotations from the cellular subset of Gene Ontology (GO) terms (Carbon et al. 2021), generated with EnTAP (Hart et al. 2020).

**OFind_Summary_Xspp** and **OFind_Full_Xspp**: The results for OrthoFinder analyses (Emms and Kelly 2019) of orthologies between *S. hermaphroditum* and various sets of proteomes from related nematodes (or, for the largest analysis set, nematodes and a nematomorph). Different numbers of proteomes in each analysis set are denoted with ‘4spp’ (four proteomes), ‘14spp’ (14 proteomes), or ‘22spp’, (22 proteomes). For each analysis, two different views of these results are given: the summary lists taxa and gene counts, while the full results give individual gene names. All proteomes used in these analyses, with their sources, are listed in Supplementary Table S7.

**OFind_Summary_4spp** and **OFind_Full_4spp**: This analysis was used to identify venom genes in *S. hermaphroditum* with maximum granularity, by orthology with known venom genes in *S. carpocapsae* or *S. feltiae*. The proteomes in this OrthoFinder analysis included: *Steinernema hermaphroditum* (from this study); *Steinernema carpocapsae*; a previous version of *Steinernema carpocapsae*; and *Steinernema feltiae*.

**OFind_Summary_16spp** and **OFind_Full_16spp**: Note that two of the ‘species’ here were different versions of *S. carpocapsae*, included to allow venom genes to be identified. This analysis was used to identify statistically significant gains and losses of gene families during the origins of *Steinernema* or *S. hermaphroditum*, and to identify strict (one-to-one) orthologies between *S. hermaphroditum* and *C. elegans*. The proteomes in this OrthoFinder analysis included: *Bursaphelenchus xylophilus*; *Caenorhabditis elegans*; *Globodera rostochiensis*; *Heterorhabditis bacteriophora*, as reannotated by Hawdon and coworkers (Vadnal et al. 2018); *Heterodera glycines*; *Panagrellus redivivus*; *Parastrongyloides trichosuri*; *Rhabditophanes diutinus* KR3021; *Steinernema hermaphroditum* (from this study); *Steinernema carpocapsae*; a previous version of *Steinernema carpocapsae*; *Steinernema feltiae*; *Steinernema glaseri*; *Steinernema monticolum*; *Steinernema scapterisci*; and *Strongyloides ratti*.

**OFind_Summary_21spp** and **OFind_Full_21spp**: This analysis was used to identify *S. hermaphroditum* genes with orthologs strongly conserved throughout the nematode phylum. The proteomes in this OrthoFinder analysis included: *Ascaris suum*; *Angiostrongylus vasorum*; *Brugia malayi*; *Bursaphelenchus xylophilus*; *Caenorhabiditis briggsae*; *Caenorhabditis elegans*; *Enterobius vermicularis*; *Globodera rostochiensis*; *Heterorhabditis bacteriophora*, as reannotated by Hawdon and coworkers (Vadnal et al. 2018); *Haemonchus contortus*; *Heterodera glycines*; *Pristionchus pacificus*; *Romanomermis culicivorax*; *Steinernema hermaphroditum* (from this study); *Steinernema carpocapsae*; *Steinernema feltiae*; *Steinernema glaseri*; *Steinernema monticolum*; *Steinernema scapterisci*; *Strongyloides ratti*; and *Trichinella spiralis*.

**Venom_orth_type**: These are venom orthology categories described in Table 2 and derived from the OrthoFinder analyses immediately below. *S. hermaphroditum* genes with orthology to a known venom gene in either *S. carpocapsae* or *S. feltiae* are annotated as follows: ‘Scarp_venom’ (orthology to an *S. carpocapsae* venom gene, but also to other *S. carpocapsae* non-venom genes), ‘Scarp_venom_only’ (orthology exclusively to *S. carpocapsae* venom genes, with no *S. carpocapsae* non-venom orthologs), ‘Sfelt_venom’ (orthology to an *S. feltiae* venom gene, but also to other *S. feltiae* non-venom genes), ‘Sfelt_venom_only’ (orthology to an *S. feltiae* venom gene, but also to other *S. feltiae* non-venom genes), and ‘Unique_Sherm’ (this gene is the only *S. hermaphroditum* member of its orthology group).

**OFind_Summary_venom** and **OFind_Full_venom**: This is a modified subset of the OrthoFinder analysis shown in **OFind_Summary_14spp** and **OFind_Full_14spp**. It is a subset because it only includes *S. hermaphroditum* genes with an orthology to either a *S. carpocapsae* venom gene or a *S. feltiae* venom gene. It is modified because venom genes are treated as a separate taxon: instead of being labeled ‘*s_carpocapsae*’ or ‘*s_feltiae*’, they are labeled ‘*s_carpocapsae.venom*’ or ‘*s_feltiae.venom*’. Marking these genes with ‘*.venom*’ allows different categories of venom gene orthology to be easily distinguished.

**GALEON_w.565.g**: This annotates genes that belong to venom gene clusters identified with GALEON (Pisarenco et al. 2024), starting with an input set of 565 broadly defined *S. hermaphroditum* venom gene orthologs.

**GALEON_w.305.g**: This annotates genes that belong to venom gene clusters identified with GALEON (Pisarenco et al. 2024), starting with an input set of more narrowly defined 305 *S. hermaphroditum* venom gene orthologs.

**Supplementary Table S3. Comparisons of Pfam protein motif frequencies.** These frequencies were computed and compared for protein-coding genes of *S. hermaphroditum* versus four other nematodes (e.g., *C. elegans*).

**Supplementary Table S4: Overrepresented protein motifs or orthology groups in venom gene clusters.** Its data columns are as follows.

**Cluster:** The specific gene cluster whose gene members were tested for statistically significant overlaps with Pfam motifs or orthology groups. These clusters were generated from an input set of 565 broadly defined *S. hermaphroditum* venom gene orthologs (and are listed in the **GALEON_w.565.g** data column of Supplementary Table S2).

**Motif or Orthology Group:** A Pfam motif or OrthoFinder orthology group whose genes were found to be significantly overrepresented in the cluster, with respect to their background frequency among all 19,428 protein-coding genes in *S. hermaphroditum*. Orthology groups are from the 16-taxon OrthoFinder analysis (annotated in data columns **OFind_Summary_16spp** and **OFind_Full_16spp** of Supplementary Table S2).

**Motif-Orth_genes:** For each Pfam motif or OrthoFinder orthology group, the number of genes encoding that motif or falling into that orthology group.

**Cluster_genes:** For each cluster, the number of venom ortholog genes falling into the cluster.

**Motif-Orth.Cluster_overlap:** The number of genes belonging both to the Pfam motif/OrthoFinder orthology group, and to the venom gene cluster.

**Exp_rand_overlap:** The number of genes that would have been randomly expected to belong both to the Pfam motif/OrthoFinder orthology group and to the venom gene cluster, given background frequencies of both categories in the total 19,428-gene set.

**Enrichment:** The ratio between **Motif-Orth.Cluster_overlap** and **Exp_rand_overlap**. The higher this ratio, the higher the overrepresentation of the Pfam motif or OrthoFinder orthology group in the venom cluster.

**q-value:** The statistical significance of this enrichment, with p-values computed by a two-tailed Fisher test, and with q-values computed from p-values to correct for multiple hypothesis testing.

**Supplementary Table S5: Overrepresented protein motifs in expanding or contracting gene families.** Statistics are given for overrepresentation of Pfam protein motifs in genes belonging to expanding or contracting gene families. Expanding or contracting gene families for two phylogenetic nodes (the origin of *Steinernema* and the splitting off of *S. hermaphroditum*; Figure 7) were extracted from the results of CAFE analysis of the 16-taxon OrthoFinder analysis (annotated in data columns **OFind_Summary_16spp** and **OFind_Full_16spp** of Supplementary Table S2), with varying thresholds of statistical significance (p ≤ 0.05, p ≤ 0.01, or p ≤ 0.001). Sheets of statistics are given for the following sets of genes.

**Stein_231fams_up.0.05:** 1,702 genes in 231 gene families expanding at the root of *Steinernema* with p ≤ 0.05.

**Stein_144fams_up.0.01:** 1,323 genes in 144 gene families expanding at the root of *Steinernema* with p ≤ 0.01.

**Stein_56fams_up.0.001:** 752 genes in 56 gene families expanding at the root of *Steinernema* with p ≤ 0.001.

**Stein_5fams_down.0.05:** three genes in five gene families contracting at the root of *Steinernema* with p ≤ 0.05.

Stein_1fams_down.0.01: one gene in one gene family contracting at the root of *Steinernema* with p ≤ 0.01.

**Sherm_159fams_up.0.05:** 1,451 genes in 159 gene families expanding at the divergence of *S. hermaphroditum* with p ≤ 0.05.

**Sherm_121fams_up.0.01:** 1,278 genes in 121 gene families expanding at the divergence of *S. hermaphroditum* with p ≤ 0.01.

**Sherm_96fams_up.0.001:** 1,127 genes in 96 gene families expanding at the divergence of *S. hermaphroditum* with p ≤ 0.001.

**Sherm_112fams_down.0.05:** 263 genes in 112 gene families contracting at the divergence of *S. hermaphroditum* with p ≤ 0.05.

**Sherm_88fams_down.0.01:** 148 genes in 88 gene families contracting at the divergence of *S. hermaphroditum* with p ≤ 0.01.

**Sherm_30fams_down.0.001:** 77 genes in 30 gene families contracting at the divergence of *S. hermaphroditum* with p ≤ 0.001.

In each sheet, the data columns are as follows.

**Motif:** A Pfam motif whose genes were found to be significantly overrepresented in the set of genes belonging to a particular group of expanding or contracting gene families.

**Motif_genes:** For each Pfam motif, the number of genes encoding that motif or falling into that orthology group.

**Family_genes:** For each set of expanding or contracting gene families, the total number of genes falling into the set.

**Motif.Family_overlap:** The number of genes belonging both to the Pfam motif and to the set of expanding or contracting gene families.

**Exp_rand_overlap:** The number of genes that would have been randomly expected to belong both to the Pfam motif and to the set of expanding or contracting gene families, given background frequencies of both categories in the total 19,428-gene set.

**Enrichment:** The ratio between **Motif.Family_overlap** and **Exp_rand_overlap**. The higher this ratio, the higher the overrepresentation of the Pfam motif in the set of expanding or contracting gene families. **q-value:** The statistical significance of this enrichment, with p-values computed by a two-tailed Fisher test, and with q-values computed from p-values to correct for multiple hypothesis testing.

**Supplementary Table S6: Software used in this study.** “Software” provides the name of each computer program or suite of computer programs; “Purpose” describes why this software was used here; “Main web site (URL) or code location” gives the primary Web site for this software; “Bioconda source (if used)” gives the bioconda web site for programs that were installed as bioconda environments; “Online documentation” gives the web site for detailed online manuals, if a given program has one. Publications and arguments for each program are cited and described in Methods.

**Supplementary Table S7: Published genomic data used in this study.** Genomic data of relevant nematodes or nematomorphs were downloaded from WormBase (Sternberg et al. 2024) or WormBase ParaSite (Howe et al. 2017) and used for transcriptomic or proteomic analyses; they include UniProt (UniProt 2023) and RefSeq (O’Leary et al. 2016) proteome databases from highly GO-annotated model organisms. Six separate spreadsheets are given for data files of genomes, repetitive DNA, proteomes, previously published RNA-seq data, protein motifs, and gene annotations in GFF format. For data files, “Species” gives the biological species described by the file, “Comments” describes the particular use(s) to which the data file was put in this study, and “URL” gives the Web source of the file.

**Supplementary File S1. Line commands for PAML analysis.** This provides detailed parameters for our use of mcmctree and codeml from PAML 4.9 to estimate species divergence times through Bayesian MCMC (Yang and Rannala 2006; Rannala and Yang 2007).

**Supplementary File S2. R script for generating input files of CAFE.** This R script was used to converted a species tree into an ultrametric tree with ape 5.8 (Paradis and Schliep 2019), and to convert an orthology group table into a table of gene counts for each orthology group and gene/species combination with data.table 1.15.4.

**Supplementary File S3: Genomic coordinates of protein-coding gene predictions in GFF3 format.**

*Sherm_braker_2024.08.14.gff3.gz*, archived at: https://osf.io/a7w9q

**Supplementary File S4: Protein sequences encoded by protein-coding genes in FASTA format.**

*Sherm_2024.08.14.pep.fa.gz*, archived at: https://osf.io/eud54

**Supplementary File S5: CDS DNA sequences encoded by protein-coding genes in FASTA format.**

*Sherm_2024.08.14.cds_dna.fa.gz*, archived at: https://osf.io/ye9qb

**Supplementary File S6: ncRNA genomic locus predictions in GFF3 format.**

*Sherm_2024.08.14.infernal_cmscan.Rfam-14.10.gff.gz*, archived at: https://osf.io/9gh5c

**Supplementary File S7: Consolidated ncRNA genomic locus predictions in GFF3 format.**

*Sherm_2024.08.14.infernal_cmscan.Rfam-14.10.plus_and_minus.merged.gff.gz*, archived at: https://osf.io/vhrgc

**Supplementary File S8: Genomic coordinates of repetitive genomic DNA elements in GFF3 format.**

*Sherm_genDNA_2024.08.14.rmask_reps.gff.gz*, archived at: https://osf.io/98wms

**Supplementary File S9: Consolidated repetitive genomic DNA locus predictions on nuclear chromosomes in GFF3 format.** *Sherm_genDNA_2024.08.14.rmask_reps.chrs_only.merged.gff.gz*, archived at: https://osf.io/pc6d3

**Supplementary File S10: DNA sequences encoded by predicted repetitive genomic DNA elements in FASTA format.** *Sherm_raven_2024.08.14_reps-families.filt1.fa.gz*, archived at: https://osf.io/28z6h

